# Novel EGFRvIII-CAR transgenic mice for rigorous preclinical studies in syngeneic mice

**DOI:** 10.1101/2021.01.31.429020

**Authors:** Pavlina Chuntova, Yafei Hou, Ryosuke Naka, Yitzhar Goretsky, Takahide Nejo, Gary Kohanbash, Tiffany Chen, Abigail L. Mende, Megan Montoya, Akane Yamamichi, Kira M. Downey, David Diebold, Jayne Skinner, Hong-Erh Liang, Bjoern Schwer, Hideho Okada

## Abstract

**Background:** Rigorous preclinical studies of chimeric antigen receptor (CAR) immunotherapy will require large quantities of consistent and high-quality CAR-transduced T (CART)-cells that can be used in syngeneic mouse glioblastoma (GBM) models. To this end, we developed a novel transgenic (Tg) mouse strain with a fully murinized CAR targeting epidermal growth factor receptor variant III (EGFRvIII).

**Methods:** We first established the murinized version of EGFRvIII-CAR and validated its function using a retroviral vector (RV) in C57BL/6J mice bearing syngeneic SB28 GBM expressing EGFRvIII. Next, we created C57BL/6J-background Tg mice carrying the anti-EGFRvIII-CAR downstream of a Lox-Stop-Lox cassette in the *Rosa26* locus. We bred these mice with CD4-Cre Tg mice to allow CAR expression on T-cells and evaluated the function of the CART-cells both *in vitro* and *in vivo*. In order to inhibit immunosuppressive myeloid cells within SB28 GBM, we also evaluated a combination approach of CART and an anti-EP4 compound (ONO-AE3-208).

**Results:** Both RV- and Tg-CART-cells demonstrated specific cytotoxic activities against SB28-EGFRvIII cells. A single intravenous infusion of EGFRvIII-CART-cells prolonged the survival of glioma-bearing mice when preceded by a lymphodepletion regimen with recurrent tumors displaying profound EGFRvIII loss. The addition of ONO-AE3-208 resulted in long-term survival in a fraction of CART-treated mice and those survivors demonstrated delayed growth of subcutaneously re-challenged both EGFRvIII^+^ and parental EGFRvIII^−^ SB28.

**Conclusion:** Our new syngeneic CAR Tg mouse model can serve as a useful tool to address clinically relevant questions and develop future immunotherapeutic strategies.

**Importance of study:** The majority of preclinical studies evaluating CART therapy for GBM have utilized xenografts implanted into immunocompromised mice. Because the successful development of these strategies will depend on the understanding of critical interactions between therapeutic cells and the endogenous immune environment, it is essential to develop a novel immunocompetent system which allows us to study these interactions in a robust and reproducible manner. To this end, we created a Tg mouse strain in which all T-cells express a murinized EGFRvIII-CAR. T-cells derived from these mice demonstrated consistent CAR expression and EGFRvIII-specific cytotoxicity while traditional transduction with a CAR vector showed batch-to-batch variability. The syngeneic system also gave us the opportunity to evaluate a combination regimen with blockade of myeloid-derived suppressor cells. The Tg-CART mice represent a novel system for robust, and reproducible preclinical investigations.

## INTRODUCTION

CART therapy represents a promising immunotherapeutic modality based on remarkable outcomes in treating hematologic malignancies.^1^ However, the development of effective CART therapies for GBM must overcome multiple barriers, including antigenic heterogeneity, CART-cell homing to GBM, and suppressive tumor microenvironment (TME), as we reviewed recently.^2–4^

Indeed, the vast majority of preclinical CART studies have utilized CAR-transduced human T-cells and xenografts in immunocompromised mice. To understand the resistance mechanisms and develop effective CART therapy strategies, including rational combinations with other immunomodulating agents, a clinically relevant and immunocompetent pre-clinical model is needed. To adequately evaluate the study questions in multiple arms, experiments will require a large quantity of GBM-bearing mice infused with consistent number and quality of syngeneic murine CART-cells. However, this aim may not be readily achieved by transducing murine T-cells with viral vectors encoding a CAR. Due to inconsistent transduction efficiency and the random genomic integration sites for the CAR transgene, the final transduced T-cell product may represent a bulk population of cells expressing varying surface levels of the CAR construct. As a result, the quantity and quality of CART-cells for mouse studies vary greatly batch-to-batch and this can limit our ability to evaluate critical questions that will facilitate improved mechanistic understanding and development of novel therapy regimens.

To overcome these challenges, in the current study, we generated a novel Tg mouse strain with *Rosa26*-targeted genetic insertion of EGFRvIII-specific^5,6^ CAR cDNA. When bred with CD4-Cre Tg mice, all T-cells derived from these Tg mice constitutively express a murinized third generation EGFRvIII-CAR, thereby serving as a continuous, reliable, and predictable source of CART-cells. We chose EGFRvIII as the CAR target in this model because the recent, first-in-human trial of i.v. administered EGFRvIII CART-cells in patients with recurrent GBM^7^ demonstrated both an encouraging safety profile and also important issues to overcome. I.v.-infused CART-cells infiltrated GBM tumors and reduced the number of EGFRvIII-positive GBM cells, but failed to demonstrate clinical benefits associated with induction of immuno-regulatory molecules, such as program cell death ligand-1 (PD-L1) in the TME.^7^

As a tumor model, we developed a C57BL/6J-background SB28 GBM cell line^8,9^ engineered to express EGFRvIII. We have shown that SB28 cells recapitulate key characteristics of human GBM, such as aggressive growth, low mutation burden, and resistance to immune checkpoint blockade therapy,^8,9^ allowing us to investigate many clinically relevant questions.

The GBM microenvironment is largely composed of immunosuppressive myeloid cells, representing an amalgam of tissue resident microglia and myeloid-derived suppressor cells (MDSCs).^10,11^ As a mechanism promoting MDSC accumulation in gliomas, we and others have shown the critical role of prostaglandin-E2 (PGE2).^8,12,13^ Therefore, we evaluated a combination of i.v. EGFRvIII-CART therapy and inhibition of immunosuppressive GBM-infiltrating myeloid cells using ONO-AE3-208, a selective inhibitor of the EP4 receptor,^14^ which is known to mediate PGE2-induced promotion of immunosuppressive myeloid cells.^15^

Our data show that the novel Tg mouse-based CAR system can be a useful and highly clinically relevant tool for the study of EGFRvIII-expressing GBM.

## MATERIALS AND METHODS

### Rosa26-targeting mCAR vector

The 3C10 antibody-derived single chain variable fragment (scFv, Suppl. Fig. 1A) was linked with murine CD8*α* hinge and transmembrane domain. The intracellular transducing domains of the murine co-stimulating factors CD28 and 4-1BB were added followed by the murine CD3(activating domain (Fig. 1A). The entire murinized construct (mCAR) sequence was codon optimized for expression efficiency and synthesized by GenScript. Binding sites for the restriction enzyme AscI were added to the 5’ and 3’ ends of the construct by PCR and used to insert the mCAR sequence into the *Rosa26*-targeting vector CTV [(Addgene #15912), Fig. 2A].

**Figure 1.**
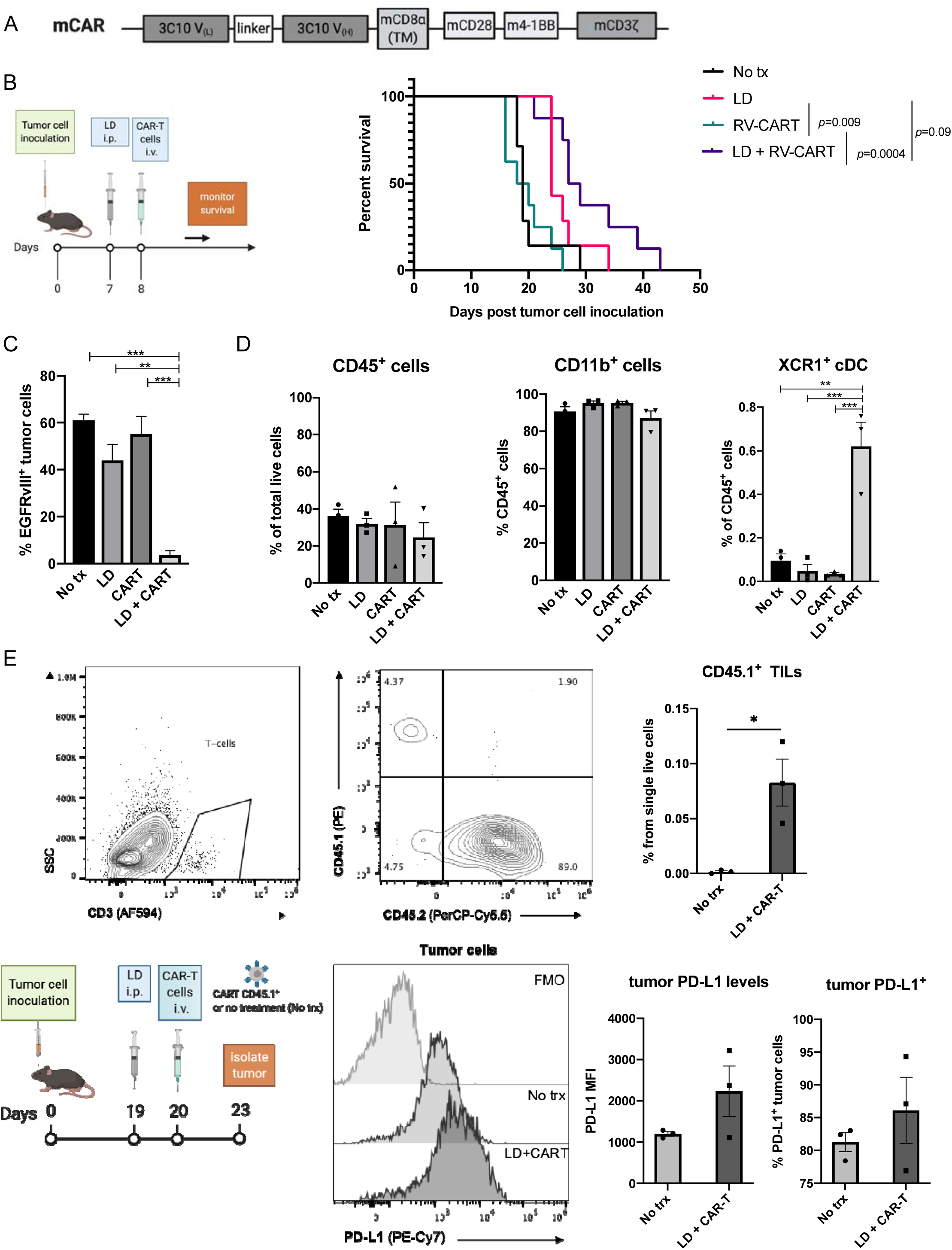
A single intravenous infusion of EGFRvIII-specific CART-cells prolongs survival of tumor-bearing mice but results in recurrence demonstrating target antigen loss. **(A)** Diagram of mCAR domains. **(B)** Schematic representation of treatment protocol and Kaplan-Meier curves demonstrating the survival. No treatment (n=7; median survival [MS] 19 days), LD (n=7; MS 24 days), RV-CART (n=8; MS 19 days), LD+RV-CART (n=8; MS 28 days). **(C)** Tumor samples from mice at endpoint were collected and analyzed by flow cytometry (FC) for surface expression of EGFRvIII. Positive staining was determined relative to an isotype control. **(D)** Summary of FC staining for myeloid populations in samples from **(C)**. **(E)** Prospective study of tumor tissues 3 days post LD+CART i.v. infusion. Tumor single-cell suspensions were analyzed by FC, and representative FC diagrams are shown.

**Fig. 2.**
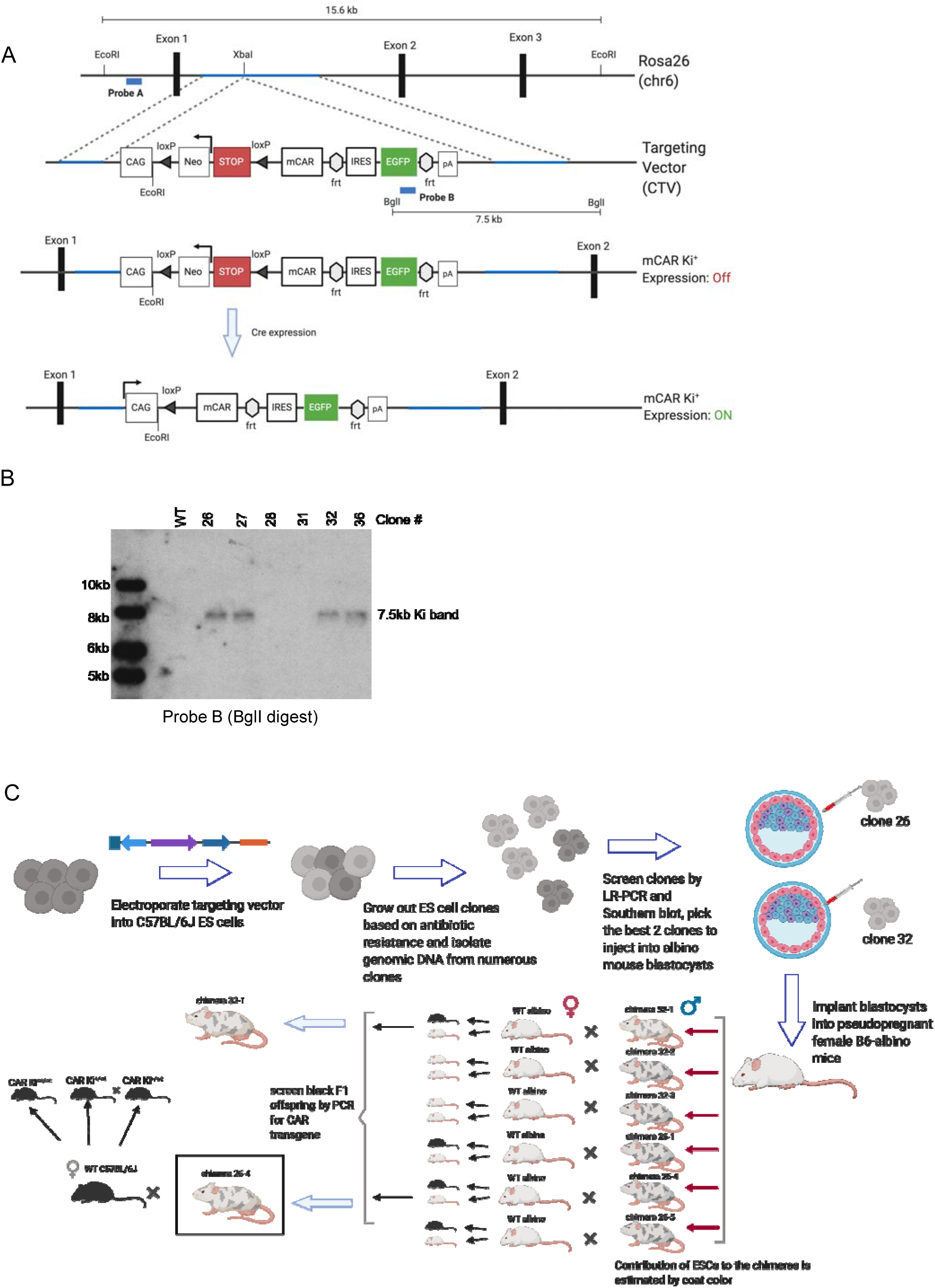
Generation of transgenic mice with the conditional EGFRvIII-CAR allele at the *Rosa26* locus. **(A)** Schematic representation of the *Rosa26* chromosome 6 locus and the location of the targeted insertion. Restriction sites for EcoRI and BglI enzymes and locations for Probes A and B are indicated. **(B)** Correct genomic targeting of the transgene at the 3' end was confirmed by Southern blotting of BglI-digest genomic DNA with Probe B. **(C)** Overview of targeting the conditional allele to the *Rosa26* locus of C57BL/6J ES cells and generation of founder mice.

### Transgenic mice

The CTV-mCAR construct was linearized using the AsiSI enzyme (New England Biolabs) and electroporated into PRXB6T (C57BL/6J) embryonic stem cells (ESC). Genomic DNA from multiple clones, resulting from G418 selection of the ESC, was screened by long-range PCR (Suppl. Fig. 2A-B) and Southern blot (Fig. 2B, Suppl. Fig. 2C-E) to confirm the intended insertion of the CTV-mCAR construct at the *Rosa26* locus. Selected ESC clones were injected into 3.5-day embryos from B6(Cg)-*Tyr^c2J^*/J (B6-albino; JAX 000058) by the Transgenic Gene Targeting Core at UCSF. A total of 62 embryos were injected and implanted into recipient B6-albino female mice. Eight chimeric male founder mice were born. The majority of tested founders (4/6) were able to transmit the transgene when bred with B6-albino females resulting in offspring with fully black coat color. The selected founder (26-4) was then bred with C57BL/6J females (JAX 000664) to establish the CAR knock-in (Ki) colony (Fig. 2C). The CAR Ki mice were bred with the commercially available CD4-Cre mouse strain (JAX 022071), and resulting offspring were genotyped by PCR for expression of both transgenes. All mice used for breeding and experiments maintained *Cre* hemizygosity to minimize any unintended consequences of the *Cre* random insertion. The primer list is available in Suppl. Table 1.

### Retroviral vectors

Standard cloning methods were used to generate the pMSCV-m3C10-IRES-GFP, pMSCV-m3C10, and pBABE-hEGFRvIII retroviral vectors. Method details are described in the Supplementary Materials and Methods.

### Retroviral supernatants

Standard packaging plasmid pCL-Eco and a gene-of-interest expressing retroviral plasmid were used to obtain retrovirus-containing medium used in all transduction assays. A detailed protocol is available in Supplementary Materials and Methods.

### Retroviral transduction of mouse T-cells

Procedure details are described in the Supplementary Materials and Methods section.

### SB28-EGFRvIII cell line

A detailed description of the cell line is available in the Supplementary Materials and Methods and Supplementary Fig. 3.

### *Real-time cytotoxicity assay* (xCelligence)

Target cells were seeded in E-Plates View 96 PET (ACEA Biosciences, no. 300600900) at a density of 5×10^3^ cells/well. Changes in electrical impedance were expressed as a dimensionless cell index (CI) value, which derives from relative impedance changes corresponding to cellular coverage of the electrode sensors, normalized to baseline impedance values with medium only. When a CI value close to 1 was reached, CART-cells were added at different effector to target (E/T) ratios ranging from 10/1 to 2.5/1. Impedance measurements were performed every 15 minutes for up to 72 hours. All conditions were performed in triplicate wells. To analyze the acquired data, CI values were exported and percentage of lysis was calculated in relation to the control cells lacking any effector CART-cells (spontaneous) using the formula [1-(experimental / spontaneous)] x100. Triton X-100 at a final concentration of 0.1% was added as maximum lysis control.

### Orthotopic glioma models

All experiments used 6–10 week-old C57BL/6J mice (Jackson Laboratory). Animals were handled in the Animal Facility at UCSF, per an Institutional Animal Care and Use Committee-approved protocol. 5×10^3^ cells were stereotactically injected through an entry site at the bregma 2 mm to the right of the sagittal suture and 3 mm below the surface of the skull of anesthetized mice using a stereotactic frame as previously described.^16^

### Isolation of tissue-infiltrating leukocytes

The procedure for the isolation of brain-infiltrating leukocytes (BILs) using Percoll (GE Healthcare Life Sciences) was previously described.^17^ To isolate tumor-infiltrating leukocytes (TILs), tumors were macro-dissected from surrounding brain tissue first.

### In vitro differentiation and analysis of mouse bone marrow

To obtain conditioned medium (CM), 0.5×10^6^ SB28-EGFRvIII cells were cultured in cRPMI. After 4 days, medium was collected, filtered through a 0.22μm sterile filter, and stored at −20°C in single-use aliquots. Levels of PGE2 in the CM were measured via the PGE2 ELISA Kit – Monoclonal (Caymann Chemical #514010) following the manufacturer’s instructions. Whole bone marrow (1×10^6^ cells) from C57BL/6J mice was cultured in 50% cRPMI/50% SB28-EGFRvIII CM either in the presence of solvent control or ONO-AE3-208 at indicated concentrations for 10 days. Fresh medium and inhibitor were added to the cultures every 4 days. Finally, adherent cells were collected and either analyzed by FC or subjected to RNA isolation using the RNeasy Mini Kit (Qiagen #74106).

### Flow cytometry

Single cell suspensions of spleens, lymph nodes, BILs, or TILs were stained with fluorescently-labeled antibodies using concentrations recommended by the manufacturers. A list of the antibodies used is available in Suppl. Table 2. Detailed description of the flow cytometry (FC) procedure is provided in Supplementary Materials and Methods.

### PCR-based genotyping

Details on the PCR-genotyping are described in Supplementary Materials and Methods.

### Reagents and drugs

Lymphodepletion: Cyclophosphamide for Injection, (Baxter, NDC 10019-955-01) was resuspended in sterile saline at 40 mg/ml immediately prior to use, and 4 mg per mouse was administered i.p. Fludarabine phosphate for Injection, (Sagent, NDC 25021-242-02, 25 mg/ml) was stored at 4°C, and 1 mg per mouse was administered i.p. EP4 inhibition *in vivo*: ONO-AE3-208 (Sigma, SML2076) was resuspended at 1 mg/ml in water, and 10 mg/kg BW were administered daily p.o.

### Statistical Analyses

Log-rank (Mantel-Cox) test by GraphPad Prism software (v8.1) was used to determine significance between Kaplan-Meier survival plots. Results from experiments with only two groups were analyzed using Student’s *t* test. For experiments with more than two groups, results were analyzed by one-way ANOVA followed by Tukey post-test. All data shown as mean ± SEM unless otherwise indicated. Symbols indicate statistical significance as follows: \**p*<0.05; \*\**p*<0.002; \*\*\**p*<0.0002.

## RESULTS

### Murinized EGFRvIII-CART-cells display EGFRvIII-specific cytotoxic effects and prolong survival of syngeneic mice bearing EGFRvIII^+^ glioma

Our first step in establishing a preclinical immunocompetent model was to evaluate the therapeutic efficacy of T-cells transduced with a retroviral vector encoding a murinized anti-EGFRvIII CAR (Fig. 1A and Suppl. Fig. 1A) in syngeneic mice bearing SB28-EGFRvIII^+^ gliomas. Similar to the parental SB28 cell line,^8,9^ the SB28-EGFRvIII tumors grow aggressively *in vivo* and result in median survival (MS) of 19 days if untreated (Fig. 1B). T-cells transduced with the murinized 3C10-CAR (CART) showed specific cytotoxicity against EGFRvIII^+^ SB28 tumor cells *in vitro* (Suppl. Fig. 4A-B). *In vivo* (Fig. 1B), CART treatment alone showed no therapeutic benefits compared to untreated mice, underscoring the need for lymphodepletion (LD) prior to systemic (i.v.) adoptive cell therapy (i.v. ACT).^18^ LD alone significantly extended the MS of tumor-bearing mice (24 days; vs No tx *p*=0.04). Combining LD and CART, however, led to the longest MS (28 days; vs CART alone *p*=0.0004). When the animals reached endpoint, we collected and examined tumors for expression of EGFRvIII (Fig. 1C). While tumors from untreated or single therapy-treated mice retained some expression of the target antigen (range 31%-70% positive; Suppl. Fig. 4D), tumor cells from combination-treated animals displayed significant loss of EGFRvIII expression (3.5 ± 1.9%, vs CART alone *p*=0.0006), strongly reminiscent of human trial observations.^7^ In the tumor microenvironment (TME), we observed that a high proportion of each tumor was composed of CD11b^+^ cells (Fig. 1D). Combination treatment did not influence this observation. However, XCR1^+^CD11b^−^ dendritic cells, which can cross-present intracellular-derived antigens,^19^ significantly increased in the animals treated with both LD and CART-cells (Fig.1D, right). In a smaller, prospective cohort, we examined the levels of PD-L1 expression 3 days after treatment with LD+CART (Fig. 1E). We detected a small population of CART-cells within the tumors which was identified in FC plots by the congenic marker CD45.1. Similarly to the observations made in clinical trial patients,^7^ PD-L1 expression on tumor cells was variable. However, 2 of 3 tumors derived from CART-infused mice showed upregulation of PD-L1 signal and an increased number of positive cells (Fig. 1E).

### Generation of transgenic mice

To overcome challenges inherent to producing large numbers of consistent and high-quality CART-cells needed for multi-arm *in vivo* studies, we generated a novel mouse strain using targeted genetic insertion of the EGFRvIII-specific CAR, which can be expressed in a tissue-specific fashion, thereby serving as a continuous, reliable, and predictable source of CART-cells. We cloned the newly created mCAR expression cassette (Fig. 1A) into the *Rosa26* targeting vector CTV (Fig. 2A) between the *Lox*-Stop-*Lox* (LSL) cassette and the *frt-*flanked IRES-EGFP sequence (i.e. Rosa-LSL-mCAR-EGFP vector). Following the electroporation of C57BL/6J ESC with the Rosa-LSL-mCAR-EGFP vector, 4 clones (26, 27, 32 and 36) had the Ki cassette in the targeted *Rosa26* locus (Fig. 2B, Suppl. Fig. 4). We selected clones #26 and #32 for further expansion and blastocyst microinjection, resulting in 8 chimeric male mice. After breeding 6 of the chimeras to albino C57BL/6J females, 4 out of 6 chimeras demonstrated germline transmission of the injected ESCs by producing offspring with black coats. Founder 26-4 was selected to establish the CAR Ki colony (summarized in Fig. 2C).

### Tg-CART-cells demonstrate EGFRvIII-specific cytotoxicity *in vitro*

To validate specific functions of the transgene, we isolated T-cells from the spleens of CAR Ki mice and transduced them with a Cre-mCherry expressing retrovirus (Fig. 3A). CAR Ki T-cells were readily transduced with the control mCherry virus (middle panels) and showed no background expression of EGFP (readout for transgene cassette expression). Specifically, after transduction with Cre, we detected expression of EGFP in both CD4^+^ and CD8^+^ T-cells (Fig. 3A bottom panels). To induce expression of the CAR in T-cells *in vivo*, we bred the CAR Ki mice to the commercially available CD4-Cre transgenic mice. CAR^*ki/wt*^; CD4-Cre^+^ (referred to as Tg mice in the rest of the studies) mice appeared anatomically healthy, without observable differences in size or weight compared to WT or CAR^*ki/wt*^; CD4-Cre^−^ controls (data not shown). Spleens from Tg mice displayed normal T-cell frequencies (Fig. 3B). As the CD4-Cre mice express Cre in all T-cells during double-positive stage of thymic development,^20^ the CAR should be expressed in both CD4^+^ and CD8^+^ mature T-cell subsets (Tg-CART-cells). In splenocytes, we observed EGFP signal in all CD4^+^ and CD8^+^ T-cells (Fig. 3C, top panel) but not NK cells, B cells, or myeloid cells (Suppl. Fig. 5A). Flow staining with anti-mouse F(ab) antibody directly detected CAR expression in splenic Tg-CART-cells. As the expression of the CAR is constitutive in all T-cells from Ki^+^ Cre^+^ animals, the absence of gross pathological phenotype suggests that the CAR reactivity is specific to the neoantigen EGFRvIII, thereby sparing the host mice from off-target toxicities.

**Figure 3.**
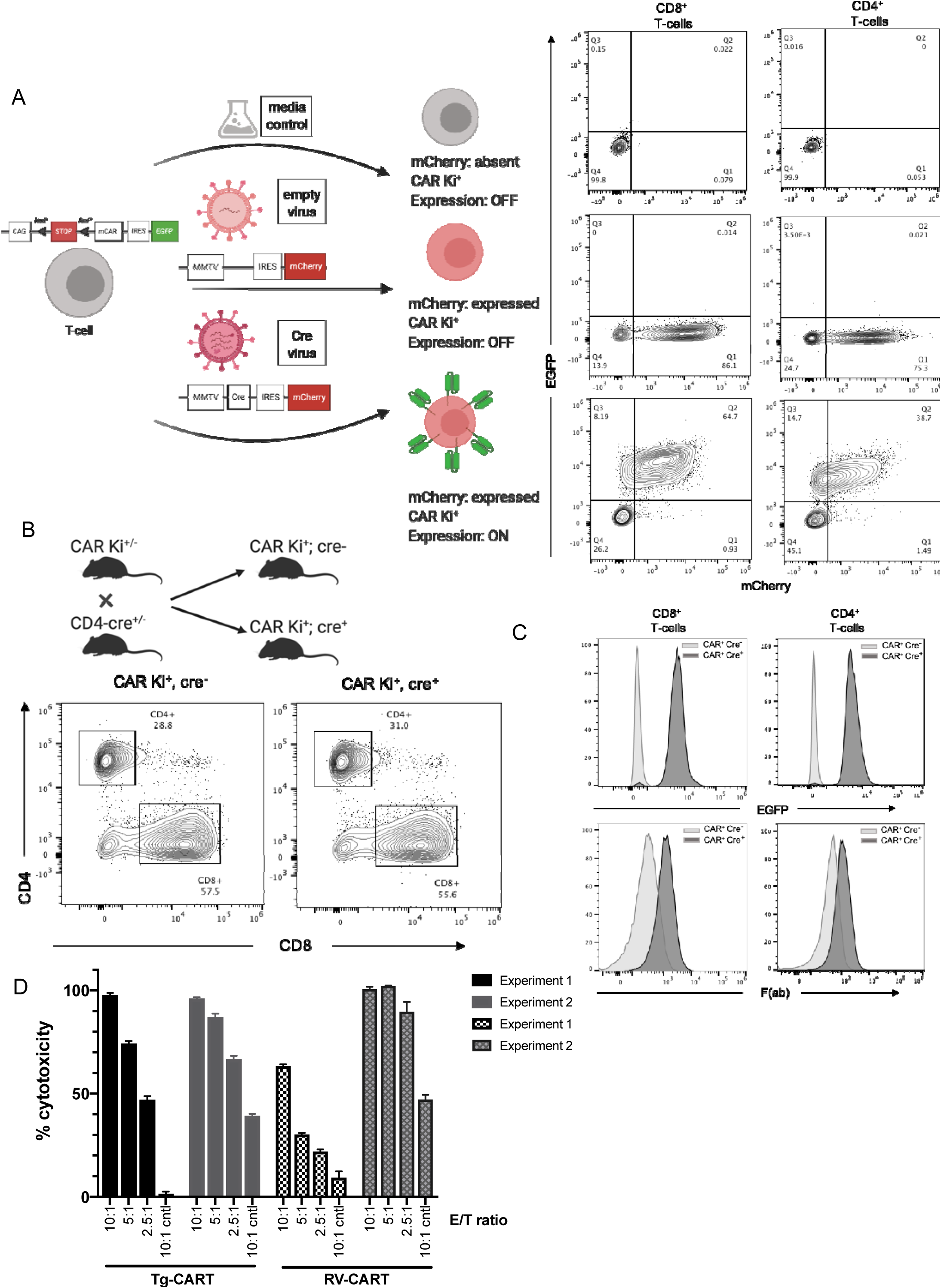
The transgene CAR is expressed following Cre recombinase expression. **(A)** Bulk CD3^+^ T-cells, isolated from the spleens of CAR Ki mice, were retrovirally transduced with a Cre-mCherry virus, a control virus expressing mCherry but not Cre (“empty virus”), or a control “no virus” media. On day 5 after transduction, CD8^+^ and CD4^+^ cells were analyzed by flow cytometry for the expression of mCherry and EGFP. **(B)** CAR Ki mice were bred with CD4-Cre mice to induce the expression of the mCAR Ki cassette *in vivo*. Representative flow plots of bulk spleen-derived CD4^+^ and CD8^+^T-cells post CD3/CD28-stimulation. **(C)** CD8^+^ and CD4^+^ T-cells from CAR Ki^+^, cre^−^ and CAR Ki^+^, cre^+^ were analyzed for the expression of EGFP and mouse F(ab). **(D)** RV- and Tg-CART-cells induce cytotoxicity against EGFRvIII^+^ SB28 cells. Sorted CD8^+^ CART-cells were co-cultured with SB28-EGFRvIII^+^ or parental SB28 (cntl) cells at different effector to target (E/T) ratios. Data from 2 separate experiments shows variability in RV-CART performance.

Next, we tested the functionality of the Tg-CART-cells by *in vitro* cytotoxicity assays. We generated CD8^+^ CART-cells through either retroviral transduction of WT C57BL/6 T-cells (RV-CART) or by isolating them from age-matched Tg mice (Tg-CART). In both groups, we activated and expanded the cells under the same conditions. Differentiation and activation markers showed no significant differences between the two groups (Suppl. Fig.6). Cytotoxicity assays against SB28-EGFRvIII^+^ cells or parental SB28 cells (cntl cells) demonstrated both RV-CART and Tg-CART-cells showed specific cytotoxicity against EGFRvIII^+^ cells but not parental SB28. (Fig. 3D). We also included control T-cells (WT incubated with medium only or isolated from CAR^*ki/wt*^; CD4-Cre^−^ mice) and saw no effect on the viability of parental or EGFRvIII^+^ cells (data not shown). It is noteworthy that cytotoxicity by Tg-CART-cells at all E/T ratios remained comparable in two separate experiments, while RV-CART-cells showed variable cytotoxicity levels between experiments (Fig. 3D). Overall, these data reveal that T-cells from the novel Tg mouse strain are able to express the EGFRvIII-CAR gene specifically and effectively upon Cre expression, thereby serving as a reliable source of EGFRvIII-specific murine CART-cells.

### Potent anti-EGFRvIII^+^ glioma effects of Tg-CART-cells *in vivo*

We then evaluated the engraftment and anti-glioma efficacy of Tg-CART-cells in comparison to RV-CART-cells *in vivo*. C57BL/6J CD45.2^+^ mice bearing day 15 SB28-EGFRvIII^+^ tumors (n=6) received LD regimen, then an i.v. infusion consisting of a 1:1 mixture of congenically marked CD45.1^+^ 2×10^6^ RV-CART-cells and 2×10^6^ EGFP^+^ CD45.2^+^ Tg-CAR-T-cells on the following day (Fig. 4A). CART-cells from both origins were sorted for CAR expression and expanded *in vitro* under identical culture conditions prior to the infusion. The flow cytometry analysis showed that the RV-CART and Tg-CART preparations consisted of 85% and 69% CD8^+^ T-cells, respectively.

**Figure 4.**
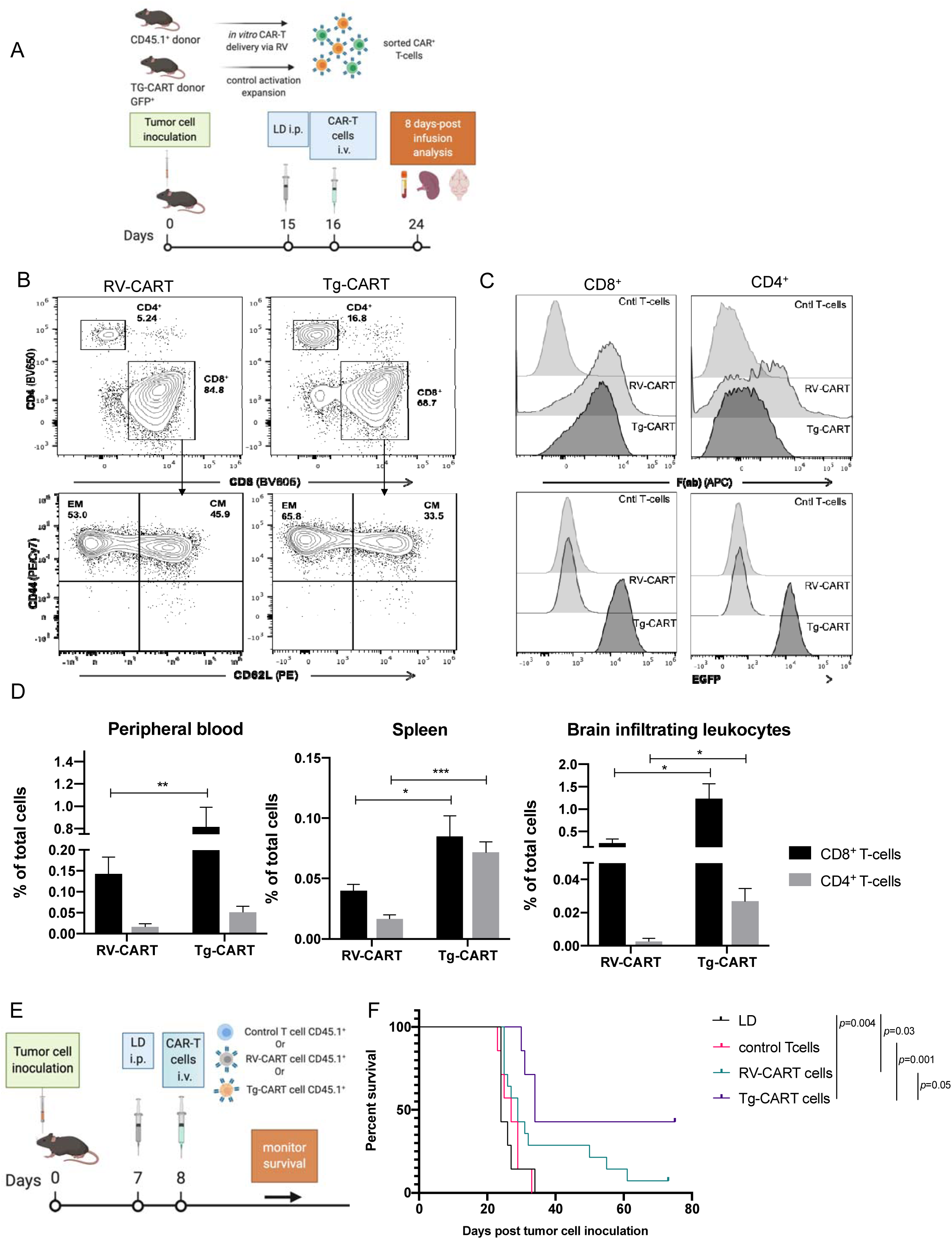
Tg-CART-cells persist better following IV infusion in tumor-bearing animals and prolong mouse survival further than RV-CART-cells. **(A)** Schematic representation of treatment protocol. **(B)** An aliquot of the infusion product was stained for CD8^+^/CD4^+^ cells, central memory (CM, CD62L^+^CD44^+^) and effector memory (EM, CD62L^−^CD44^+^) in each CART preparation. **(C)** F(ab) staining (top panels) denoting CAR expression and EGFP levels (bottom panels) in each CART preparation. Both T-cell preparations were activated *in vitro* and expanded under the same conditions. **(D)**Tissues were analyzed by flow cytometry for the presence of CAR-T-cells, where RV-CART-cells were CD45.1^+^EGFP^−^ and Tg-CART-cells were CD45.2^+^EGFP^+^. Percentages were calculated from all single viable cells acquired from each sample. Bars represent the mean of 6 biological replicates. **(E)** Schematic representation of treatment protocol. **(F)** Kaplan-Meier curves: LD group (MS = 24 days, n=7), control T-cells (MS = 27 days, n=7), RV-CART-cells (MS = 29 days, n=14), and Tg-CART-cells (MS = 34 days, n=7).

In the CD8^+^ compartment, the two populations displayed similar amounts of central memory (CM, CD62L^+^CD44^+^) T-cells (RV- vs Tg-CART: 46% vs 34%) and effector memory (EM, CD62L^−^CD44^+^) T-cell (Fig. 4B lower panels). CD4^+^ cells from both groups also showed comparable levels of CM vs EM differentiation (Suppl. Fig. 7A). CD8^+^ T-cells showed similar CAR expression, as shown by levels of F(ab) staining (Fig. 4C), whereas RV-derived CD4^+^ CART-cells appeared to contain a subpopulation of cells which showed higher levels of CAR expression compared with Tg-CART-cells (Fig. 4C, top right panel). EGFP expression, as expected, was only present in Tg-CART-cells. To examine the cells’ ability to traffic to and engraft in different tissues, we analyzed lymphoid and non-lymphoid tissues from animals sacrificed 8 days after the i.v. ACT (Fig. 4D). CD8^+^ Tg-CART-cells were detected at higher levels in the blood and spleen compared to CD8^+^ RV-CART-cells. No significant difference in the levels of CD4^+^ cells was seen in the blood, while CD4^+^ Tg-CART-cells were ~3 times more abundant in the spleen than their RV-CART counterparts. Among leukocytes isolated from the tumor-bearing brain hemispheres, both CD8^+^ and CD4^+^ Tg-CART-cells were observed at higher levels than the RV-CART-cells (Fig. 4D). These data indicated that despite going through identical activation and culture protocols and showing similar activation profile at time of infusion, Tg-CART-cells survived better in the periphery and persisted longer at the tumor site than RV-CART-cells.

Next, to evaluate the therapeutic efficacy of Tg-CART-cells, mice bearing day 7 SB28-EGFRvIII^+^ gliomas received LD and i.v. ACT of either CAR^−^ T-cells (control), RV-CART-cells, or Tg-CART-cells on the following day (Fig. 4E). While RV-CART-cells prolonged survival of treated mice (29 days; vs LD *p*=0.03) including one long-term survivor, treatment with Tg-CART-cells resulted in a longer median survival (34 days; vs LD *p*=0.004) and long-term tumor control in 3/7 treated animals (Fig. 4F), although the difference did not reach a statistical significance in comparing the RV-CART and Tg-CART cohorts (*p*=0.05).

### Inhibition of PGE2 pathway enhances the efficacy of anti-EGFRvIII CART therapy

Similarly to human GBM, a large component of the immune TME in SB28-EGFRvIII tumors consists of MDSC (Suppl. Fig. 8A and B) which were not significantly altered by the CART infusion (Suppl. Fig. 8C). Furthermore, SB28-EGFRvIII cells produced high levels of PGE2 in the culture supernatant (Fig. 5A). While PGE2 acts by binding to its receptors EP2 and EP4 on myeloid cells,^21^ mouse bone-marrow derived CD11b^+^ cells cultured in the presence of SB28 conditioned media (CM) demonstrated the expression of EP4, but only background levels of EP2 (Fig. 5B). A small molecule antagonist of EP4, ONO-AE3-208,^14^ significantly decreased the SB28-EGFRvIII CM induced expression of Arginase I (ArgI), a known mediator of MDSC-mediated immunosuppression,^22^ at both mRNA (Fig. 5C) and protein levels (Fig. D). Furthermore, treatment with ONO-AE3-208 effectively decreased the ratio of CD206^+^MHCII^−^/CD206^−^MHCII^+^ CM-cultured CD11b^+^ cells (Suppl. Fig9A-B), suggesting a shift towards an anti-tumor myeloid cell phenotype.^23^These data point to the PGE2/EP4 signaling axis as a major contributor to MDSC development in SB28 tumors and led us to test the therapeutic efficacy of ONO-AE3-208 *in vivo*.

**Figure 5.**
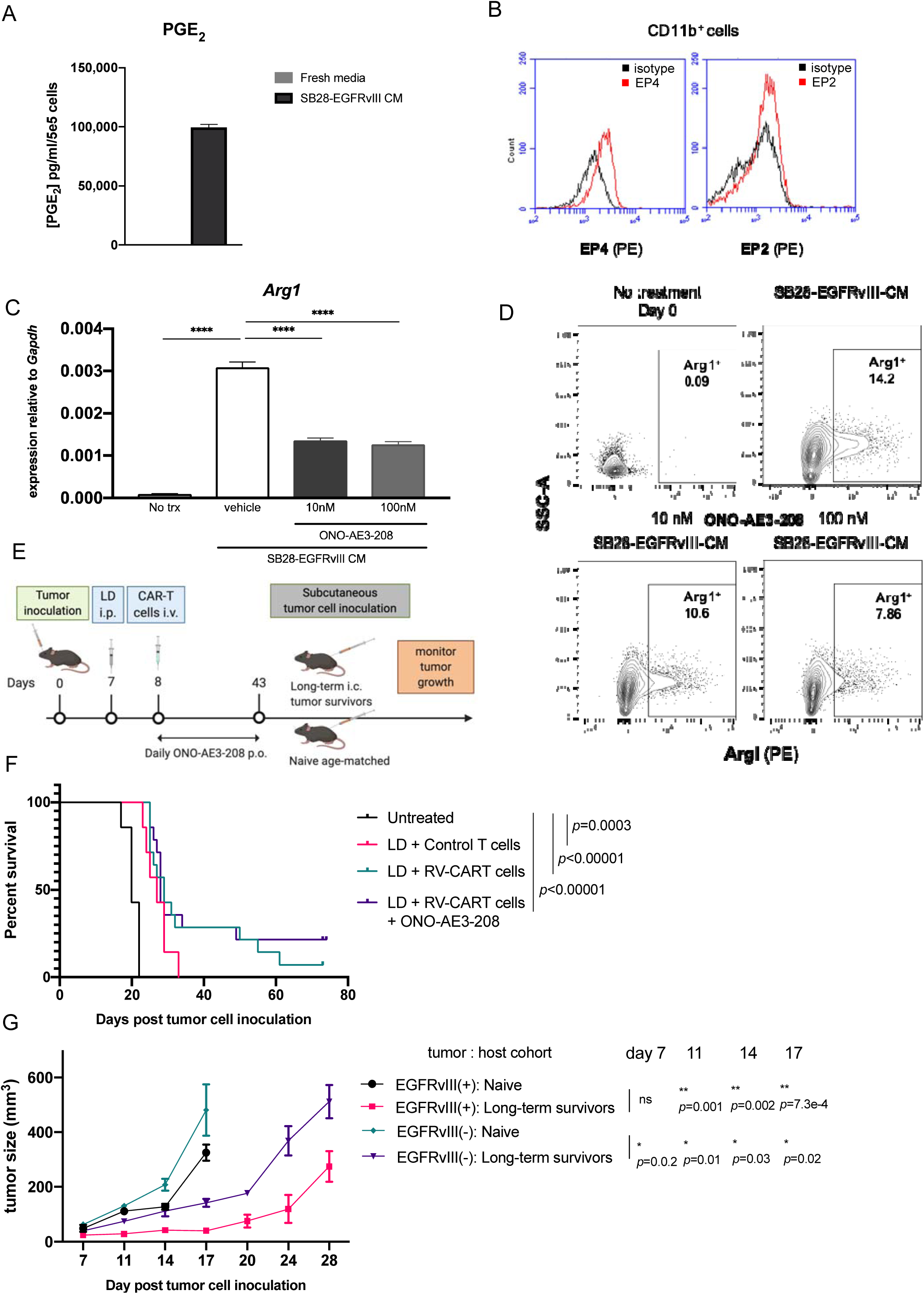
Inhibition of the PGE2 pathway improves the efficacy of anti-EGFRvIII CAR-T therapy. **(A)** PGE2 production by SB28-EGFRvIII glioma cells by specific ELISA **(B)** Flow cytometry analysis of EP2 and EP4 on bone-marrow-derived CD11b^+^ cells following 6-day culture in the presence of SB28-CM; the black histogram representing unstained cells, and red histogram represents cells stained with anti-EP2 or anti-EP4 antibody. **(C)** and **(D)** ONO-AE3-208 inhibits *Arg1* mRNA **(C)** and protein **(D)** expression in CD11b^+^ cells exposed to SB28-EGFRvIII CM. **(E)**Schematic representation of treatment protocol for evaluation of the combination regimen with CART therapy and ONO-AE3-208. Mice receiving the CART therapy were further randomized to receive daily p.o. treatment of either vehicle (n=7) or 10mg/kg ONO-AE3-208 (n=7) for up to 35 consecutive days. **(F)**Kaplan-Meier curves demonstrating overall survival. Untreated (MS = 20 days, n=7), LD+Control T-cells (MS = 27 days,a n=7), LD+RV-CART-cells (MS = 29 days, n=14), and LD+RV-CART-cells+ONO-AE3-208 (MS = 28 days, n=14) **(G)** Long-term survivors from the LD+CAR-T+ONO-AE3 group received subcutaneous (s.c.) injections of 4×10^5^ parental SB28 cells (EGFRvIII^−^, left flank) and 4×10^5^ SB28-EGFRvIII cells (EGFRvIII^+^, right flank). Tumor size (in mm^3^) was measured every 3 days. Naïve, age-matched C57Bl6/J mice received identical s.c. injections as controls.

Mice bearing day 7 SB28-EGFRvIII tumors were stratified to receive no treatment (n=7) or LD (n=35) (Fig. 5E). The following day, we administered LD-treated mice with an i.v. infusion of either control-(n=7) or RV-CART-cells (n=28). In addition, we began daily p.o. treatment of the RV-CART-treated cohort with either vehicle (n=14) or 10mg/kg ONO-AE3-208 (n=14) for up to 35 days. As in earlier experiments, both LD (MS=27 days, vs Untreated *p*=0.0003) and LD in combination with RV-CART (MS=29 days, vs Untreated p<0.0001) extended the survival of tumor-bearing mice significantly. Although ONO-AE3-208 did not extend the median survival of CART-treated mice (MS=28 days), we observed extended survival in 3/14 (21%) animals treated as far out as day 73, whereas only 1/14 (7%) mouse treated with RV-CART alone remained tumor-free on day 73 (Fig. 5F). On day 74, we re-challenged the ONO-treated long-term survivors with injections of parental SB28 cells (EGFRvIII^−^; left flank) and SB28-EGFRvIII^+^ cells (right flank). As a control group, we also injected age-matched naïve mice (n=3) and measured tumor volumes every 3 days (Fig. 5E). EGFRvIII^+^ tumors in naïve mice grew significantly faster than in the treated mice (Fig. 5G). Furthermore, the growth of the parental SB28 cells was also inhibited in the treated mice compared with the naïve cohort (*p*<0.05 at all timepoints, Fig. 5G). These data suggest that the treatment regimen of RV-CART + ONO-AE3-208 in mice bearing EGFRvIII^+^ gliomas induced a memory response to antigens present in both parental and EGFRvIII^+^ cells; i.e. “antigen spreading” following the initial targeting of EGFRvIII.

## DISCUSSION

In this study, we developed a novel Tg mouse strain with a *Rosa26*-targeted insertion of a murinized anti-EGFRvIII CAR. Along with the syngeneic SB28-EGFRvIII glioma cells, this system should serve as a consistent and readily available source of CART-cells for clinically relevant studies in syngeneic C57BL/6 mice. To our knowledge, this is the first development of *Rosa26*-targeted CAR-Tg mice. The field has greatly appreciated the availability of mice with transgenic T-cell receptors (TCR), such as ones against gp100 (Pmel)^24^ or ovalbumin (OT-1^25^ and OT-2^26^). The consistent quality and availability of large number of T-cells derived from these mice allow us to perform experiments with adequate scientific rigor and to compare results between experiments within a project and across multiple studies that test different strategies. As there have been few CAR-based preclinical studies of GBM using syngeneic models,^27–30^ our EGFRvIII CAR Tg-mice will likely be highly valuable for future preclinical studies in the neuro-oncology field. Because EGFRvIII is a tumor-specific antigen, we hypothesized that despite the 90% homology between human and mouse EGFR, constitutive expression of the CAR on T-cells in CAR Ki CD4-Cre mice would not cause any adverse effects. Our data show that the new transgenic mice are viable, fertile, and display no aberrant phenotype. Additionally, the Tg-CART-cells offer the possibility to easily access therapeutic cells that can be easily tracked *in vivo* using EGFP or a congenic marker like CD45.1^+^.

With regards to CAR-Tg mice, the Kershaw lab created a Tg mouse strain expressing a CAR against Her2 under the control of the pan-hematopoietic promoter, Vav, and evaluated the ability of these mice to respond to Her2 expressing tumors.^31^ Subsequently, Kershaw and colleagues generated dual-specific T-cells expressing both a Her2-specific CAR and a gp100-specific TCR and demonstrated potent anti-tumor effects against a variety of Her2^+^ tumors.^32^ While the Her2-CAR Tg mice have random chromosomal insertions of the CAR transgene^33^, our model is the first to target the CAR insertion to a chromosomal safe harbor locus.

The *Rosa26*-targeted feature may have contributed to the observed superior *in vivo* functions of Tg-CART-cells compared with RV-CART-cells. In recent studies with human T-cells, it was also shown that genomic locus-targeted insertion of a CAR^34^ or TCR^35^ transgene confers superior efficacy. To eliminate any bias introduced by different T-cell subpopulations in our *in vitro* studies, both T-cell groups consisted of only CAR^+^ CD8^+^ cells. We noted that Tg-CART-cells displayed uniform but lower CAR expression levels based on F(ab) staining, while RV-CART-cells showed varied expression levels. Increased uniformity of CAR expression on Tg-CART-cells likely owes to a single copy of the cassette in the *Rosa26* locus. Although the optimal level of CAR density is unknown, too high a density may result in early CART-cell exhaustion due to ligand-independent tonic signaling by the CD3ζ due to CAR molecule clustering.^36^ Our mCAR vector design includes the 4-1BB co-stimulation molecule which has been shown to reduce the exhaustion effects of persistent CAR signaling.^36^ Together, our data suggest that CAR expression from a single allele in a known locus may provide CART-cells with enhanced efficacy and support the development of locus-targeted engineering of human T-cells.^34,35^

We and others have studied and reviewed the important contributions of the myeloid components within the glioma microenvironment.^37–41^ The use of fully syngeneic, immunocompetent preclinical models is necessary for studying the immunosuppressive effects of glioma-associated myeloid cells on both the endogenous anti-tumor response and any therapeutic infusions. Our model allowed us to test the hypothesis that inclusion of the EP4 antagonist ONO-AE3-208 would enhance the therapeutic efficacy of CART-cells. Our *in vitro* data suggest that the inhibitor prevents MDSC development and suppresses their ability to produce ArgI, thereby partially relieving the immunosuppression within the TME. Our re-challenge experiments demonstrated that the anti-tumor response in long-term surviving mice extended not only to EGFRvIII^+^ tumors but also to SB28 parental tumors lacking the CART antigen. These data suggest that, aided by the action of ONO-AE3-208, our i.v. infused CART-cell therapy was able to initiate an endogenous anti-tumor response against non-EGFRvIII antigens.

GBM possesses substantial degree of inter- and intratumor heterogeneity, which may contribute to the failure of antigen-targeted immunotherapeutic approaches.^42^ In our data, despite the sorting of EGFRvIII^+^ cells before intracerebral inoculation (Suppl. Fig. 3C), the mice which eventually succumbed to tumor burden in the LD+CART cohort recurred with tumors that lacked EGFRvIII expression. This antigen loss strongly suggests tumor immunoediting as a result of CART-cell activity. To better understand the effects of CART treatment on the GBM microenvironment in our model, we evaluated the expression of immune-related genes in the tumor via Nanostring on day 4 following Tg-CART or control T-cell infusion (Suppl. Fig. 10A-B). As differential expression analyses showed a great level of inter-individual heterogeneity (Suppl. Fig. 10C-D), further studies are warranted to validate the observed trends and understand mechanisms for efficacy and resistance. Therefore, our model presents a valuable opportunity to develop strategies to overcome antigen loss and immune escape following CART treatment in a clinically relevant tumor system.

As successful immunotherapy for primary GBM faces numerous challenges,^43^ our new Tg mouse strain allows access to high number of EGFRvIII-CART-cells with a reproducible phenotype within a flexible preclinical *in vivo* system. Coupled with our syngeneic tumor model, the Tg-CART mice will be a valuable new tool to address issues including antigen immune escape, the highly immunosuppressive TME, and whether approaches combining separate therapies can improve outcomes for GBM patients.

## Authorship

*Concept and design*: PC and HO; *Methodology:* PC, YH, RN, TN, HEL, BS, and HO; *Data acquisition:* PC, YH, RN, YG, TN, GK, TC, ALM, MM, AY, KMD, DD, JS, and HEL; *Analysis and data interpretation*: PC, YH, RN, YG, TN, GK, BS, and HO; *Manuscript preparation*: PC, ALM, KMD, BS, and HO. *Study supervision:* HO

## Acknowledgements

Tohru Kotani, Payal Watchmaker, Bindu Hegde, Ryan Gilbert, Davide Ruggero and Ruggero lab members; UCSF Core Facilities: Laboratory for Cell Analysis, Genome Analysis Core, Preclinical Therapeutics Core, and Gladstone Transgenic Gene-Targeting Core. All experimental diagrams were created with BioRender.

