## Supplemental Materials and Methods for "Novel EGFRvIII-CAR transgenic mice for rigorous preclinical studies in syngeneic mice"

**Supplementary Materials and Methods**

***Retroviral vectors***

The codon optimized mCAR sequence was inserted into the pMSCV-IRES-GFP retroviral vector (Addgene # 20672) using EcoRI and XhoI restriction enzymes. For generating RV-CART-cells for use *in vivo,* IRES-GFP was excised from the vector using the SalI and XhoI restriction enzymes. The full-length sequence of hEGFRvIII was PCR cloned from pT3.5/CMV-EGFRvIII (Addgene # 20280)^1^ and inserted into pBABE-puro (Addgene # 1764)^2^ using the SnaBI and SalI restriction enzymes. All plasmids were sequenced and stock DNA was prepared using the EndoFree Plasmid Kit (Qiagen, #12362).

***Retroviral supernatants***

The viral vector of interest and the packaging plasmid pCL-Eco were transfected into 70% confluent HEK293T cells using Fugene HD (Promega, E2311) following the manufacturer’s protocol. 18 hours later, fresh medium was added, and the retrovirus-containing medium was collected 48 and 72 hours after transfection. The viral supernatant was passed through a 45-µm filter and stored at -80$^{\circ}$C in single-use aliquots.

***Retroviral transduction of mouse T-cells***

CD3^+^ T-cells were isolated from the spleens of 6-10-week-old C57Bl/6J mice using the MojoSort Mouse CD3 T Cell Isolation kit (Biolegend, 480024) following the manufacturer’s instructions. Isolated T-cells were resuspended at 1x10^6^ cells/ml in complete RPMI [cRPMI: RPMI 1640 media with 10% FBS, 1% Penicillin-Streptomycin (Gibco, 15070063), 1% HEPES (Gibco, 15630080), 1% Glutamax (Gibco, 35050061), 1% non-essential amino acids (Gibco, 11140076), 1% sodium pyruvate (Gibco, 11360070), 0.5mM 2-Mercaptoethanol (Gibco, 21985023)] supplemented with hIL-2 (30U/ml, NIH) and 50ng/ml mIL-15 (Peprotech, 21015) and incubated with CD3/CD28 mouse T-activator beads (Gibco, 11453D) at a 1:1 ratio for 2 days. Meanwhile, wells in a 24-well non-tissue culture treated plate were coated with 20µg/ml RetroNectin (Takara Bio USA, T100B) and stored at 4$^{\circ}$C overnight. On the day of transduction, RetroNectin-coated plates were loaded with 0.5ml per well viral supernatant and centrifuged at 2,000*g* for 30 min at 32$^{\circ}$C. At the same time, T-cells were harvested, magnetized to remove activator beads, washed, and resuspended at 1-2x10^6^ cells/ml in viral supernatant and polybrene (Sigma, final concentration 8µg/ml). Aliquots of 0.5-1x10^6^ T-cells were added to each well pre-loaded with virus during the previous step, and plate was centrifuged at 1,000*g* for 1 hour at 32$^{\circ}$C. Immediately after, half of each well’s volume was carefully removed, and fresh media and cytokines were added (30U/ml hIL-2, 50ng/ml mIL-15). Cell density was monitored daily and maintained in fresh media and cytokines at 0.5-1x10^6^ cells/ml.

***The SB28-EGFRvIII cell line***

The previously described murine glioma cell line SB28 was retrovirally transduced with human EGFRvIII and expanded under puromycin selection. Resulting cells were flow sorted based on a fluorescently labeled antibody (1µg/10^6^ cells) and cultured in cRPMI media containing 10µg/µl puromycin. Surface expression of EGFRvIII was regularly assayed and confirmed via flow cytometry. Both parental and EGFRvIII-expressing tumor cells were tested routinely and confirmed mycoplasma-negative.

To reduce the immunogenicity of the EGFRvIII-transduced SB28 cells, *β2-microgloblin* (*β_2_m)* was knocked out. Two separate sgRNA sequences targeting Exon 1 of the *β_2_m* gene were used, as well as a control scrambled sgRNA. Modified sgRNAs were synthesized by Synthego (CRISPR Revolution sgRNA EZ Kit) and were stored as 100µM single use aliquots at -20°C. Sequences are listed in Suppl. Table 3 and were obtained from Das et al.^3^ Cas9-NLS protein was recombinantly produced and purified (QB3 Macrolab). The protein was stored at 40µM in 20mM HEPES-KOH, pH 7.5, 150mM KCl, 10% glycerol, 1mM TCEP in single use aliquots at -80°C and was brought to room temperature and diluted to 20µM immediately prior to use. Reconstituted sgRNA were diluted to 30µM and slowly mixed with Cas9 (9:1 sgRNA to Cas9 molar ratio). The mixture was incubated at 37°C for 15min to form a ribonucleoprotein (RNP) complex. The RNPs were electroporated into the tumor cells immediately after forming. SB28-EGFRvIII tumor cells were harvested by trypsinization and immediately prior to electroporation resuspended in the Lonza electroporation buffer SF using 100µl solution per 0.5x10^6^ cells. Cell suspension was gently mixed with RNP solution, transferred to a Lonza 100µl nucleocuvette (Cat No V4XC-2024), and electroporated using the Lonza 4D-Nucleofector X-unit with pulse code EN150. Immediately after electroporation, 500µl of pre-warmed cRPMI media were added to each cuvette, and cells were allowed to rest for 15min at 37°C. After 15min, cells were transferred to a 6-well tissue culture plate containing pre-aliquoted 1.5ml of cRPMI for a final volume of 2ml per well. Cells were cultured under normal conditions until confluent. Bulk population was assayed by flow cytometry for surface expression of MHC-I (IA^b^), and negative cells were sorted to generate an MHC-I knock-out (K/O) SB28-EGFRvIII cell line.

The SB28-EGFRvIII MHC-I K/O cells were injected orthotopically into C57BlL6J mice, as described previously. When tumor-bearing mice reached the prespecified endpoint for humane euthanasia, the tumors were harvested, dissociated into single-cell suspension, and plated out in 100mm dishes in cRPMI containing puromycin. The media was changed twice weekly until glioma cells grew as monolayers. Cells emerging from multiple tumors were assayed by flow cytometry for the surface expression of EGFRvIII, and the cell line which showed highest antigen retention was selected for all further *in vivo* studies.

***In vitro cytotoxicity measurements***

The CytoTox 96 nonradioactive cytotoxicity assay (Promega, G1780) shown in Suppl. Fig.3B was performed according to the manufacturer’s protocol. Target tumor cells (5x10^3^/well) were plated in U-bottom 96-well plates with various effector/target (E/T) ratios in 200 μl media for 24 hrs. Fifty µl of supernatant was then transferred to a flat-bottom 96-well assay plate containing 50 μl CytoTox 96 Reagent and incubated for 30 min at room temperature. Fifty µl stop solution was then added to each well, and plates were analyzed at 490 nm on a Synergy2 microplate reader (Biotek). Percentage cytotoxicity was calculated as [(experimental – effector spontaneous – target spontaneous)/(target maximum – target spontaneous)] x100.

***Flow cytometry***

Single cell suspensions (0.5-1x10^6^ cells) of spleens, lymph nodes, BILs, or TILs were pre-incubated with blocking solution (BioLegend, 156604). After 10 min, a cocktail of fluorophore-conjugated antibodies resuspended in 50 µl FC buffer (1X PBS, 0.5% FBS, 2 mM EDTA) was added directly to each tube, and samples were incubated at 4°C for 20 min in the dark. Samples were washed with excess buffer and resuspended for analysis. In the case of staining for CAR, biotinylated goat anti-mouse F(ab) (Abcam, ab5886; 1 µg/10^6^ cells) was added first to blocking solution and incubated at 4°C for 20 min. Samples were then washed twice. Fluorophore-conjugated antibody cocktail, with the addition of fluorescently-labeled streptavidin, was added as described above. Samples were analyzed using BD Accuri C6 (BD Biosciences) or the Invitrogen Attune NxT (Thermo Fisher Scientific) flow cytometers. Tumor cells were sorted using SONY SH800 (Sony Biotechnology).

***Nanostring data normalization and analysis***

Background correction was performed with the Background Thresholding option in nSolver software (v4.0), whereby a threshold count value was set to 11, and all counts which fell below that value were adjusted to match it. All the subsequent computational analyses were performed using R (v4.0.3). For the subsequent differential expression (DE) analysis, the factors of unwanted variation were estimated based on the housekeeping genes and implemented into the design matrix, according to the RUVSeq workflow.^4^ DE analysis was performed with the aforementioned design matrix using the DESeq2 R package (v1.30.0).^5^ For multiple testing, p-values were adjusted with the Benjamini-Hochberg method.

***PCR-based genotyping***

Detection of the mCAR insertion was performed by PCR using the following protocol: 94ºC 3 min, 35 cycles of 94°C 45 s, 60°C 45 s, 72°C 45 s, final extension 72°C 5 min. Reaction contained all 3 primers listed in Suppl. Table 1 at the following final concentrations: Rosa_5bisF at 1.33μM, Rosa3ArmR at 0.67μM, and PreCAG2-Rev at 0.67μM. WT allele band product is 257bp; mCAR transgene band is 326bp.

Detection of the CD4-cre transgene was performed by PCR using the following protocol: 94ºC 2 min, 10 cycles of 94°C 20 s, 65°C 15 s (decreasing by 0.5ºC per cycle), 68°C 10 s, 32 cycles of 94°C 15 s, 60°C 15 s, 72°C 10 s, final extension 72°C 2 min. Reaction contained all 3 primers listed in Suppl. Table 1 at the following final concentrations: CD4creF at 0.8μM, CD4creWT at 0.4μM, and CD4creMUT at 0.4μM. WT product band is 153bp; CD4-cre MUT band is 336bp.

Detection of EGFP excision following cross with FlpO mouse strain was achieved through PCR using the following protocol: 94ºC 3 min, 35 cycles of 94°C 45 s, 64°C 30 s, 72°C 75 s, final extension 72°C 5 min. Reaction contained both primers listed in Suppl. Table 1 was a final concentration of 0.5µM.

Genomic DNA for all reactions was isolated from tail tissue samples using DirectPCR Lysis Reagent (Tail) and Proteinase K according to manufacturer’s instructions (Viagen Biotech). Results were analyzed on a 0.005% SYBR Safe DNA Gel Stain, 1.5% agarose gel.
