## Supplemental Figures for "Novel EGFRvIII-CAR transgenic mice for rigorous preclinical studies in syngeneic mice"

### Supplemental Figure 1

**A** 3C10-mCAR sequence

```
              10        20        30        40        50        60        70        80        90       100
       *    *    *    *    *    *    *    *    *    *    *    *    *    *    *    *    *    *    *    *
     1 AscI
     |
   1 GGCGCGCCGCCACCATGGCAAGCCCTCTCACAAGGTTCCTCTCCCTGAATCTGCTCCTCATGGGCGAATCAATCATCCTGGGTAGTGGCGAAGCTGGATC 100
     G  A  P  P  P  W  Q  A  L  S  Q  G  S  S  P  *  I  C  S  S  W  A  N  Q  S  S  W  V  V  A  K  L  D  P

             110       120       130       140       150       160       170       180       190       200
       *    *    *    *    *    *    *    *    *    *    *    *    *    *    *    *    *    *    *    *
 101 CGAAATTCAGCTCCAGCAGTCTGGCGCAGAACTGGTGAAACCCGGAGCCTCAGTCAAGCTGAGCTGTACCGGCTCCGGCTTCAACATCGAGGACTACTAT 200
      E  I  Q  L  Q  Q  S  G  A  E  L  V  K  P  G  A  S  V  K  L  S  C  T  G  S  G  F  N  I  E  D  Y  Y
      1
      >>>>>>>>>>>>>>>>>>>>>>>>>>>>>>>>>>>>>>>>>>>>>>>>>>>>>>>>>>>>>>>>>>>>>>>>>>>>>>>>>>>>>>>>>>>>>>>>
      3C10 V(l)
       K  F  S  S  S  S  L  A  Q  N  W  *  N  P  E  P  Q  S  S  *  A  V  P  A  P  A  S  T  S  R  T  T  I

             210       220       230       240       250       260       270       280       290       300
       *    *    *    *    *    *    *    *    *    *    *    *    *    *    *    *    *    *    *    *
 201 ATTCACTGGGTGAAACAGCGGACTGAACAGGGTCTGGAGTGGATCGGCCGCATTGACCCTGAAAACGATGAGACCAAGTACGGACCAATCTTTCAGGGGA 300
     I  H  W  V  K  Q  R  T  E  Q  G  L  E  W  I  G  R  I  D  P  E  N  D  E  T  K  Y  G  P  I  F  Q  G  R
     34
     >>>>>>>>>>>>>>>>>>>>>>>>>>>>>>>>>>>>>>>>>>>>>>>>>>>>>>>>>>>>>>>>>>>>>>>>>>>>>>>>>>>>>>>>>>>>>>>>>
     3C10 V(l)
      F  T  G  *  N  S  G  L  N  R  V  W  S  G  S  A  A  L  T  L  K  T  M  R  P  S  T  D  Q  S  F  R  G

             310       320       330       340       350       360       370       380       390       400
       *    *    *    *    *    *    *    *    *    *    *    *    *    *    *    *    *    *    *    *
 301 GGGCCACCATTACAGCTGACACCAGCTCCAATACAGTGTATCTCCAGCTGTCTAGTCTGACTAGCGAGGATACCGCAGTGTACTATTGTGCCTTCAGAGG 400
       A  T  I  T  A  D  T  S  S  N  T  V  Y  L  Q  L  S  S  L  T  S  E  D  T  A  V  Y  Y  C  A  F  R  G
     68
     >>>>>>>>>>>>>>>>>>>>>>>>>>>>>>>>>>>>>>>>>>>>>>>>>>>>>>>>>>>>>>>>>>>>>>>>>>>>>>>>>>>>>>>>>>>>>>>>>
     3C10 V(l)
     G  P  P  L  Q  L  T  P  A  P  I  Q  C  I  S  S  C  L  V  *  L  A  R  I  P  Q  C  T  I  V  P  S  E  E

             410       420       430       440       450       460       470       480       490       500
       *    *    *    *    *    *    *    *    *    *    *    *    *    *    *    *    *    *    *    *
 401 AGGAGTCTACTGGGGCCCCGGCACCACCCTGACAGTGTCAAGCGGGGGTGGCGGATCAGGAGGTGGAGGAAGCGGAGGTGGAGGCAGCCACATGGATGTG 500
      G  V  Y  W  G  P  G  T  T  L  T  V  S  S                                                H  M  D  V
     101                                                                                      1
     >>>>>>>>>>>>>>>>>>>>>>>>>>>>>>>>>>>>>>>>>>>>>>>>>>>>>>>>>>>>>>>>>>>>>>>>>>>>>>>>>>>>>>>>>>>>>>>>>
     3C10 V(l)                              linker                                            3C10 V(h)
       E  S  T  G  A  P  A  P  P  *  Q  C  Q  A  G  V  A  D  Q  E  V  E  E  A  E  V  E  A  A  T  W  M  W

             510       520       530       540       550       560       570       580       590       600
       *    *    *    *    *    *    *    *    *    *    *    *    *    *    *    *    *    *    *    *
 501 GTCATGACACAGTCTCCCCTCACTCTGAGTGTCGCTATCGGCCAGAGTGCATCAATTAGCTGCAAATCCTCTCAGTCTCTGCTCGACAGTGATGGGAAGA 600
     V  M  T  Q  S  P  L  T  L  S  V  A  I  G  Q  S  A  S  I  S  C  K  S  S  Q  S  L  L  D  S  D  G  K  T
     5
     >>>>>>>>>>>>>>>>>>>>>>>>>>>>>>>>>>>>>>>>>>>>>>>>>>>>>>>>>>>>>>>>>>>>>>>>>>>>>>>>>>>>>>>>>>>>>>>>>
     3C10 V(h)
      S  *  H  S  L  P  S  L  *  V  S  L  S  A  R  V  H  Q  L  A  A  N  P  L  S  L  C  S  T  V  M  G  R

             610       620       630       640       650       660       670       680       690       700
       *    *    *    *    *    *    *    *    *    *    *    *    *    *    *    *    *    *    *    *
 601 CCTACCTGAATTGGCTGCTCCAGCGACCCGGCCAGAGTCCTAAAAGACTCATCAGCCTGGTGTCCAAGCTGGACTCTGGAGTCCCTGATAGGTTTACAGG 700
       Y  L  N  W  L  L  Q  R  P  G  Q  S  P  K  R  L  I  S  L  V  S  K  L  D  S  G  V  P  D  R  F  T  G
     39
     >>>>>>>>>>>>>>>>>>>>>>>>>>>>>>>>>>>>>>>>>>>>>>>>>>>>>>>>>>>>>>>>>>>>>>>>>>>>>>>>>>>>>>>>>>>>>>>>>
     3C10 V(h)
     P  T  *  I  G  C  S  S  D  P  A  R  V  L  K  D  S  S  A  W  C  P  S  W  T  L  E  S  L  I  G  L  Q  A

             710       720       730       740       750       760       770       780       790       800
       *    *    *    *    *    *    *    *    *    *    *    *    *    *    *    *    *    *    *    *
 701 CTCCGGATCTGGGACAGACTTCACTCTCCGCATCTCCCGAGTGGAAGCCGAGGATCTGGGCATCTACTATTGCTGGCAGGGCACCCACTTCCCAGGGACA 800
      S  G  S  G  T  D  F  T  L  R  I  S  R  V  E  A  E  D  L  G  I  Y  Y  C  W  Q  G  T  H  F  P  G  T
     72
     >>>>>>>>>>>>>>>>>>>>>>>>>>>>>>>>>>>>>>>>>>>>>>>>>>>>>>>>>>>>>>>>>>>>>>>>>>>>>>>>>>>>>>>>>>>>>>>>>
     3C10 V(h)
       P  D  L  G  Q  T  S  L  S  A  S  P  E  W  K  P  R  I  W  A  S  T  I  A  G  R  A  P  T  S  Q  G  H

             810       820       830       840       850       860       870       880       890       900
       *    *    *    *    *    *    *    *    *    *    *    *    *    *    *    *    *    *    *    *
 801 TTTGGGGGTGGCACTAAGCTGGAGATCAAGGCGGCCGCAACTACCACAAAGCCCGTGCTGCGGACACCCTCACCTGTCCATCCTACTGGAACCTCTCAGC 900
     F  G  G  G  T  K  L  E  I  K              T  T  T  K  P  V  L  R  T  P  S  P  V  H  P  T  G  T  S  Q  P
     105                                       1
     >>>>>>>>>>>>>>>>>>>>>>>>>>>>>>            >>>>>>>>>>>>>>>>>>>>>>>>>>>>>>>>>>>>>>>>>>>>>>>>>>>>>>>>>>
     3C10 V(h)                                 mCD8a
      L  G  V  A  L  S  W  R  S  R  R  P  Q  L  P  Q  S  P  C  C  G  H  P  H  L  S  I  L  L  E  P  L  S

             910       920       930       940       950       960       970       980       990      1000
       *    *    *    *    *    *    *    *    *    *    *    *    *    *    *    *    *    *    *    *
 901 CACAGCGACCAGAGGACTGTCGGCCTCGCGGGAGCGTGAAGGGTACTGGCCTGGACTTCGCTTGTGACATCTACATCTGGGCTCCACTGGCAGGCATCTG 1000
       Q  R  P  E  D  C  R  P  R  G  S  V  K  G  T  G  L  D  F  A  C  D  I  Y  I  W  A  P  L  A  G  I  C
     22
     >>>>>>>>>>>>>>>>>>>>>>>>>>>>>>>>>>>>>>>>>>>>>>>>>>>>>>>>>>>>>>>>>>>>>>>>>>>>>>>>>>>>>>>>>>>>>>>>>
     mCD8a
      H  S  D  Q  R  T  V  G  L  A  G  A  *  R  V  L  A  W  T  S  L  V  T  S  T  S  G  L  H  W  Q  A  S  A

             1010      1020      1030      1040      1050      1060      1070      1080      1090      1100
       *    *    *    *    *    *    *    *    *    *    *    *    *    *    *    *    *    *    *    *
1001 CGTCGCACTGCTCCTGTCCCTCATCATTACCCTGATTTGCTATCACCGATCTAGGAACAGTAGGAGAAATCGGCTCCTCCAGGTCACTACCATGAACATG 1100
      V  A  L  L  L  S  L  I  I  T  L  I  C  Y  H  R  S  R
      55
      >>>>>>>>>>>>>>>>>>>>>>>>>>>>>>>>>>>>>>>>>>>>>>>>>>>>>>>>>>>>>>>>>>>>>>>>>>>>>>>>>>>>>>>>>>>>>>>>
      mCD8a                                         mCD28
       S  H  C  S  C  P  S  S  L  P  *  F  A  I  T  D  L  G  T  V  G  E  I  G  S  S  R  S  L  P  *  T  *

             1110      1120      1130      1140      1150      1160      1170      1180      1190      1200
       *    *    *    *    *    *    *    *    *    *    *    *    *    *    *    *    *    *    *    *
1101 ACCCCTCGGCGCCCAGGACTGACAAGGAAGCCCTACCAGCCTTATGCCCCAGCTAGAGACTTTGCCGCTTACAGGCCCAAATGGATCAGAAAGAAATTCC 1200
      >>>>>>>>>>>>>>>>>>>>>>>>>>>>>>>>>>>>>>>>>>>>>>>>>>>>>>>>>>>>>>>>>>>>>>>>>>>>>>>>>>>>>>>>>>>>>>>>
      mCD28                                                                    m4-1BB
       P  L  G  A  Q  D  *  Q  G  S  P  T  S  L  M  P  Q  L  E  T  L  P  L  T  G  P  N  G  S  E  R  N  S

             1210      1220      1230      1240      1250      1260      1270      1280      1290      1300
       *    *    *    *    *    *    *    *    *    *    *    *    *    *    *    *    *    *    *    *
1201 CACATATCTTCAAGCAGCCCTTCAAGAAAACAACTGGCGCAGCCCAGGAGGAAGATGCTTGCTCCTGTAGGTGCCCACAGGAGGAAGAGGGAGGAGGTGG 1300
      >>>>>>>>>>>>>>>>>>>>>>>>>>>>>>>>>>>>>>>>>>>>>>>>>>>>>>>>>>>>>>>>>>>>>>>>>>>>>>>>>>>>>>>>>>>>>>>>
      m4-1BB
      H  I  S  S  S  S  P  S  R  K  Q  L  A  Q  P  R  R  K  M  L  A  P  V  G  A  H  R  R  K  R  E  E  V  E

             1310      1320      1330      1340      1350      1360      1370      1380      1390      1400
       *    *    *    *    *    *    *    *    *    *    *    *    *    *    *    *    *    *    *    *
1301 AGGATATGAACTGCGGGCCAAGTTCAGTCGCTCAGCTGAGACTGCTGCAAACCTCCAGGACCCTAATCAGCTGTACAACGAGCTGAACCTGGGACGAAGG 1400
      >>>>>>>>>>>>>>>>>>>>>>>>>>>>>>>>>>>>>>>>>>>>>>>>>>>>>>>>>>>>>>>>>>>>>>>>>>>>>>>>>>>>>>>>>>>>>>>>
      m4-1BB      mCD3z
        D  M  N  C  G  P  S  S  V  A  Q  L  R  L  L  Q  T  S  R  T  L  I  S  C  T  T  S  *  T  W  D  E  G

             1410      1420      1430      1440      1450      1460      1470      1480      1490      1500
       *    *    *    *    *    *    *    *    *    *    *    *    *    *    *    *    *    *    *    *
1401 GAAGAGTATGACGTGCTGGAGAAGAAGAGGGCCAGGGACCCCGAGATGGGAGGCAAGCAGCAGAGACGGCGCAACCCCCAGGAAGGCGTCTACAACGCCC 1500
      >>>>>>>>>>>>>>>>>>>>>>>>>>>>>>>>>>>>>>>>>>>>>>>>>>>>>>>>>>>>>>>>>>>>>>>>>>>>>>>>>>>>>>>>>>>>>>>>
      mCD3z
       K  S  M  T  C  W  R  R  R  G  P  G  T  P  R  W  E  A  S  S  R  D  G  A  T  P  R  K  A  S  T  T  P

             1510      1520      1530      1540      1550      1560      1570      1580      1590      1600
       *    *    *    *    *    *    *    *    *    *    *    *    *    *    *    *    *    *    *    *
1501 TCCAGAAGGACAAAATGGCAGAAGCCTATAGCGAGATCGGTACAAAAGGCGAGCGAAGGAGAGGAAAGGGGCACGATGGACTCTACCAGGGGCTGTCCAC 1600
      >>>>>>>>>>>>>>>>>>>>>>>>>>>>>>>>>>>>>>>>>>>>>>>>>>>>>>>>>>>>>>>>>>>>>>>>>>>>>>>>>>>>>>
      mCD3z
      S  R  R  T  K  W  Q  K  P  I  A  R  S  V  Q  K  A  S  E  G  E  E  R  G  T  M  D  S  T  R  G  C  P  Q

             1610      1620      1630      1640      1650      1660
       *    *    *    *    *    *    *    *    *    *    *    *
                                                          1656 AscI
                                                          |
1601 AGCCACTAAGGACACCTACGACGCACTCCACATGCAGACCCTCGCTCCAAGATGAGGCGCGCC 1663
        P  L  R  T  P  T  T  H  S  T  C  R  P  S  L  Q  D  E  A  R
```

### Supplemental Figure Legends

**Suppl. Figure 1. m3C10-CAR cDNA sequence. (A)** Annotated sequence of the mCAR cDNA, including the added AscI restriction sites.

**Supplemental Figure 2**

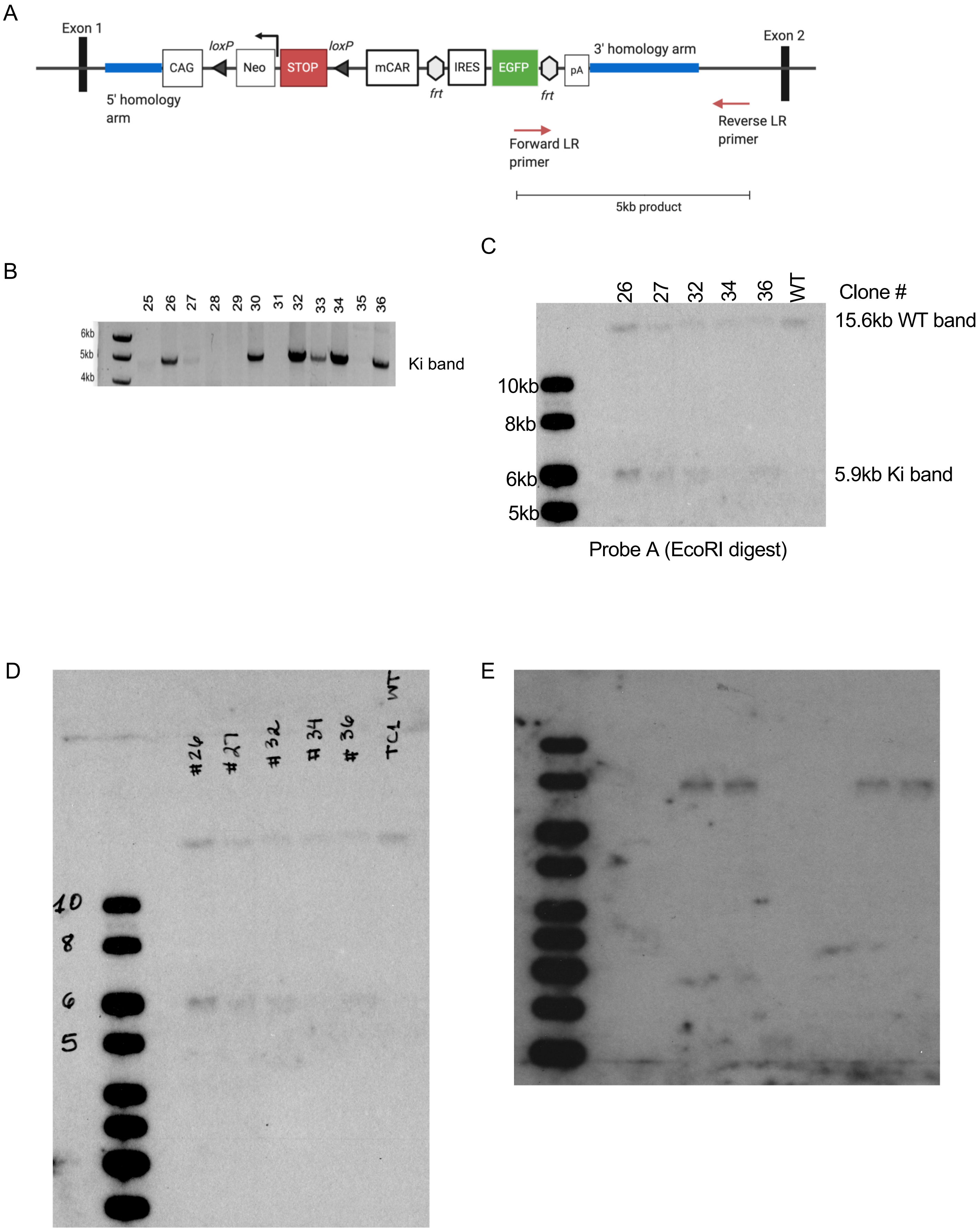

**Suppl. Figure 2. Generation of CAR Ki Tg mice (A)** Location of LR-PCR primers used to screen ESC clones. Red arrows indicate where LR-PCR primers bind within the transgene cassette (forward primer) and downstream of the 3’ homology recombination arm (reverse primer), generating a 5kb product only when the CAR sequence is targeted correctly within the *Rosa26* locus. **(B)** A 5-kb LR-PCR product identifies correctly-targeted ES colonies. **(C)** Southern blot of genomic DNA digested with EcoRI. Probe A hybridizes to two DNA fragments of different size, the 15.6-kb band indicating the presence of a WT *Rosa26* allele, while the smaller 5.9-kb band is generated by correct insertion of the knockin cassette into the *Rosa26* locus. ESC clones 26, 27, 32, and 36 showed faint but definitive bands at 15.6-kb (WT allele) and at 5.9-kb (knockin allele) indicating the clones were hemizygous at the *Rosa26* locus. **(D)** Unedited, full-length blot shown in **(C)**. **(E)** Unedited, full length Southern blot image shown Fig. 2B.

**Supplemental Figure 3**

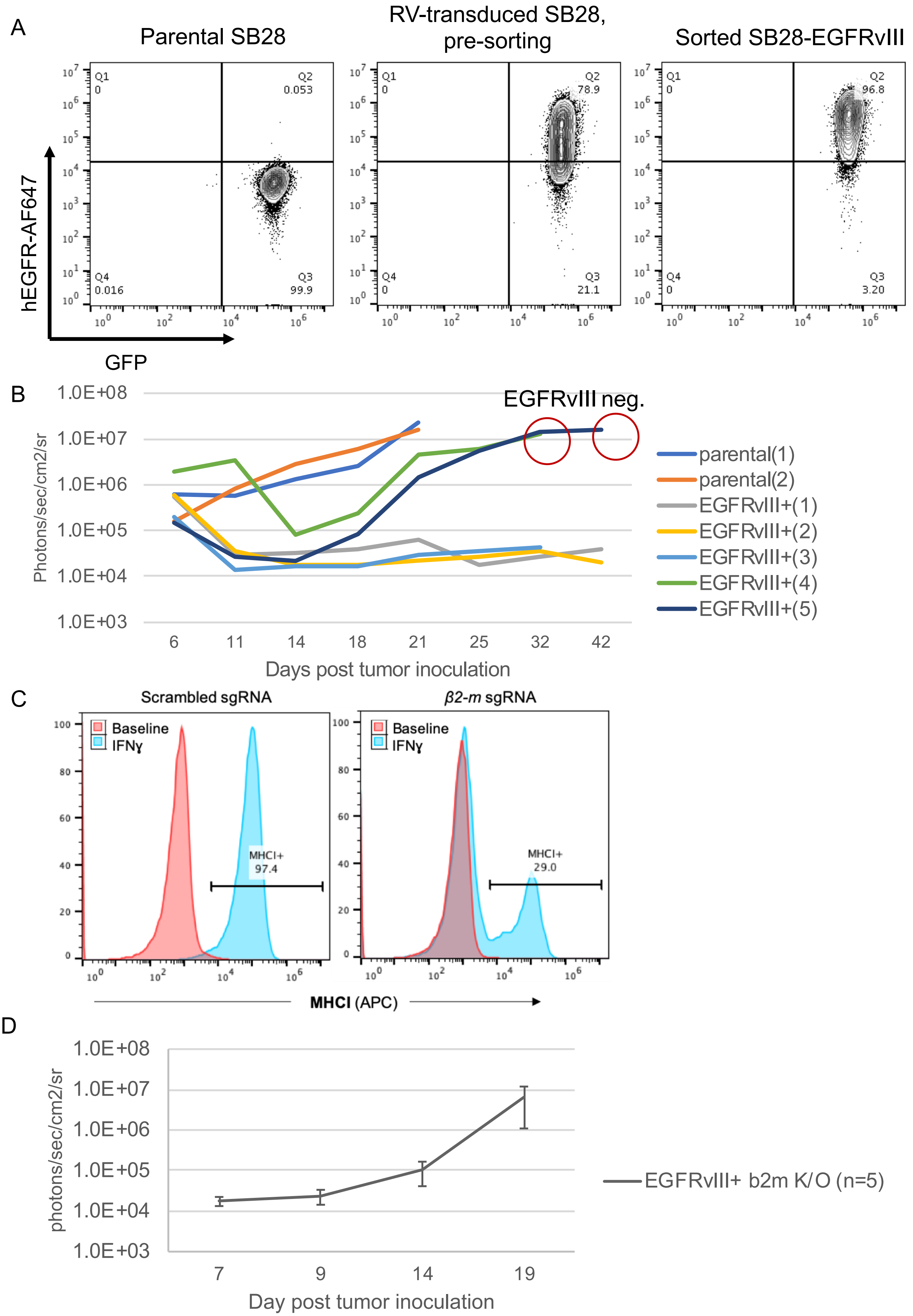

**Suppl. Figure 3. Generation of the SB28-EGFRvIII cell line (A)** SB28 cells were transduced with hEGFRvIII-expressing retrovirus and flow sorted. **(B)** Tumor growth kinetics measured by BLI signal. Animals injected with parental SB28 (n=2) reached endpoint at day 28, while those injected with SB28-EGFRvIII cells either rejected the tumors (n=3) or succumbed to EGFRvIII(-) tumors (n=2). **(C)** Knock-out (K/O) of MHC-I in SB28-EGFRvIII cells via CRISPR-mediated deletion within *β-2m*. Tumor cells do not express MHC-I at baseline. After overnight treatment with IFNγ, control cells upregulate MHC-I, while K/O cells do not. **(D)** Tumor growth kinetics. MHC-I K/O-sorted cells from **(C)** grow reproducibly in mice.

**Supplemental Figure 4**

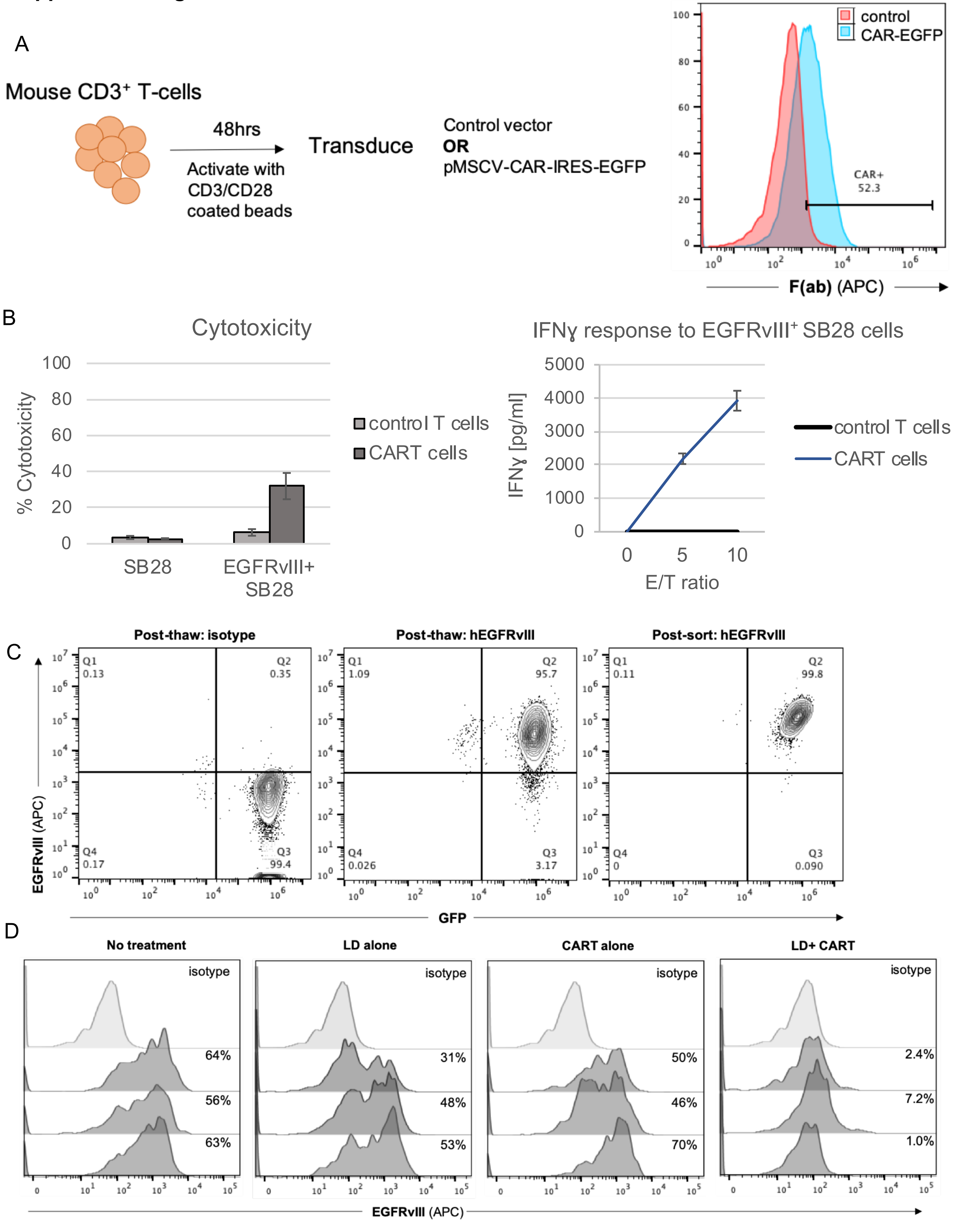


Gated on single, live, GFP+

**Suppl. Figure 4. RV-CART cells demonstrate cytotoxicity *in vitro* and *in vivo* (A)** RV-CAR transduction**.** Bulk $CD3^+$ murine T cells transduced with mCAR retrovirus stain positive for mouse F(ab) expression by FC **(B)** CART cells from **(A)** kill SB28-EGFRvIII cells and produce IFNγ *in vitro* **(C)** Representative flow plots of early-passage SB28-EGFRvIII cells before and after sorting in preparation of *in vivo* inoculation. **(D)** FC histograms of EGFRvIII levels in SB28-EGFRvIII tumors treated in Fig. 1B.

**Supplemental Figure 5**

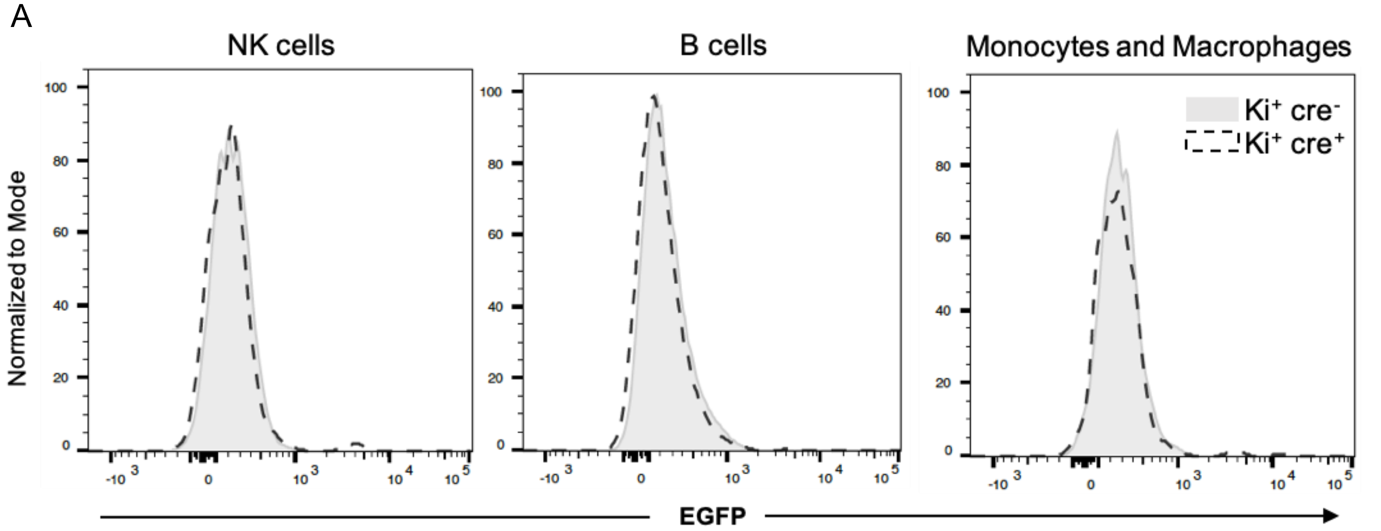


Gated on:
live, singlets, CD11b+, NK1.1+

Gated on:
live, singlets, B220+, CD3Neg

Gated on:
live, singlets, CD11b+, NK1.1Neg; CD11cNeg; Ly6GNeg

**Suppl. Figure 5. Characterization of Tg-CART mice (A)** NK cells, B cells, and myeloid cells from Tg-CAR mice do not express the CAR. Splenocytes from Ki+ cre- and Ki+ cre+ mice were stained and analyzed by FC for expression of CAR (evaluated by EGFP levels).

**Supplemental Figure 6**

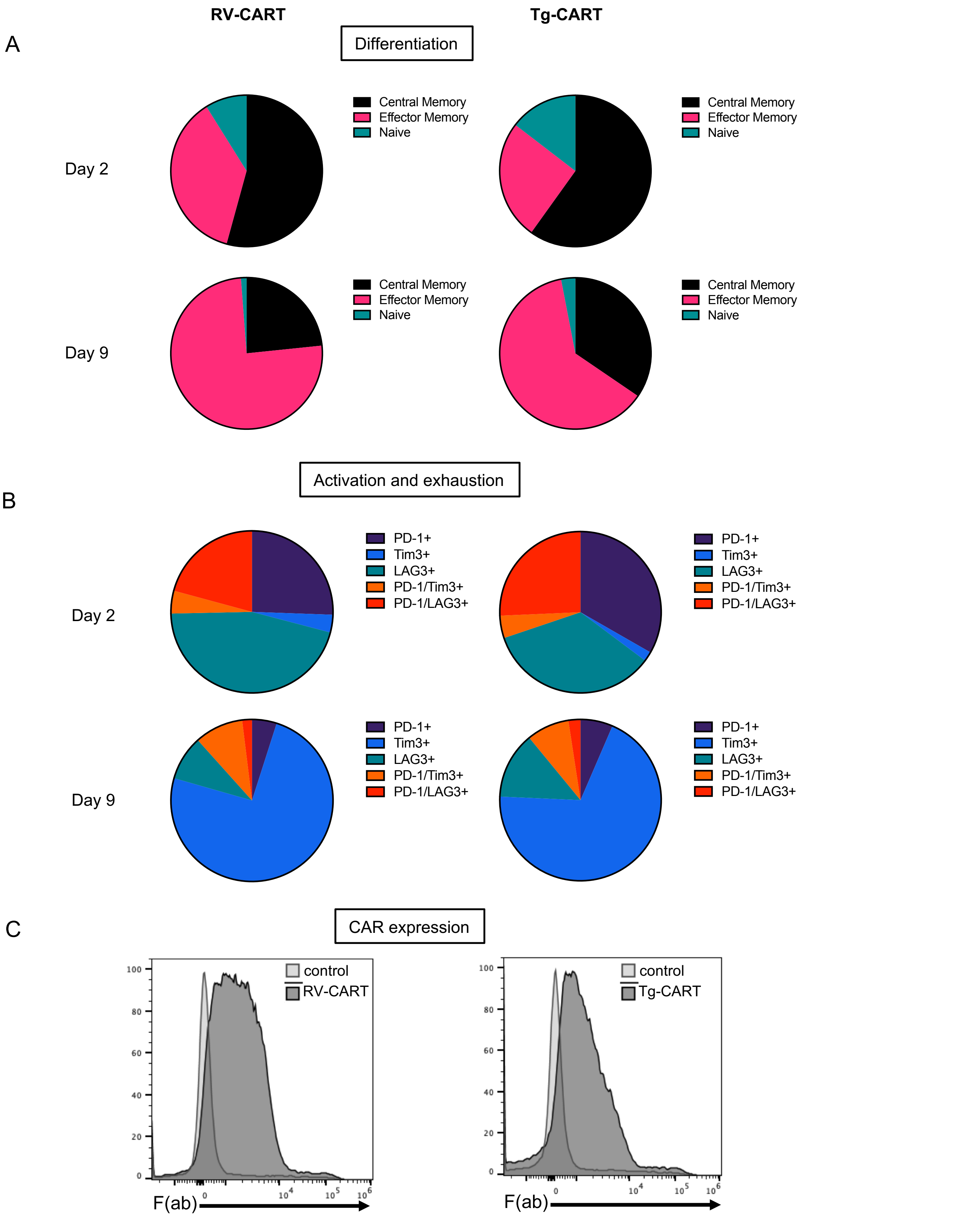

**Suppl. Figure 6. RV- and Tg-CART cells display similar differentiation and activation profiles. (A)** $CD8^+$ T cells from both populations were stained and analyzed by FC for the expression of CD62L and CD44. Percentages of naïve ($CD62L^+$, $CD44^-$), central memory (CM, $CD62L^+$, $CD44^+$), and effector memory (EM, $CD62L^-$, $CD44^+$) populations on days 2 and 9 post transduction were shown. **(B-C)** Cells from (A) were analyzed for the expression of activation/exhaustion markers (B) and surface levels of CAR following a F(ab)-based sort (C).

### Supplemental Figure 7

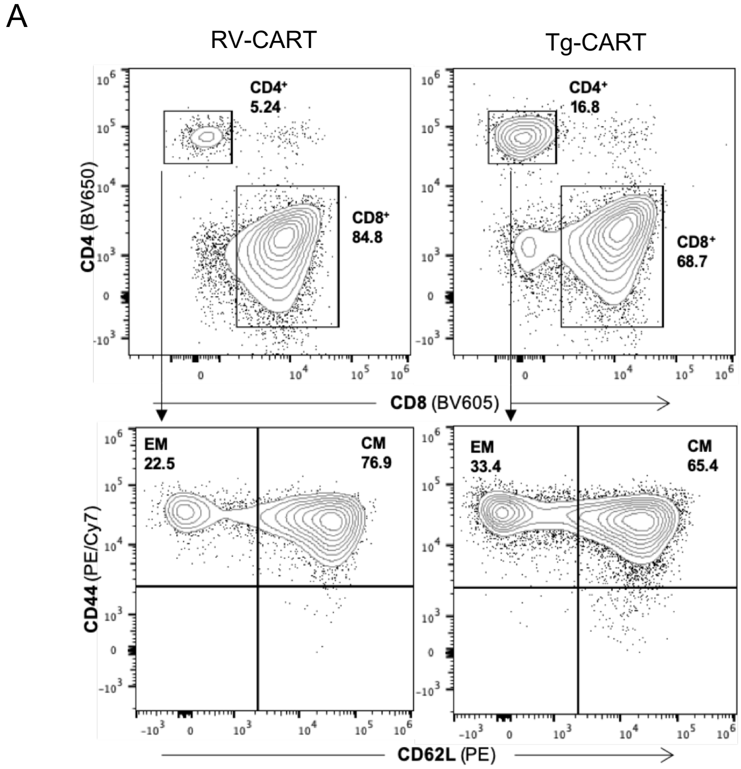

**Suppl. Figure 7. CD4$^{+}$ RV- and Tg-CART cells show similar differentiation profiles (A)** CD4$^{+}$ T cells used in the infusion product from Fig. 4A have similar percentages of CM and EM cells regardless of origin.

**Supplemental Figure 8**

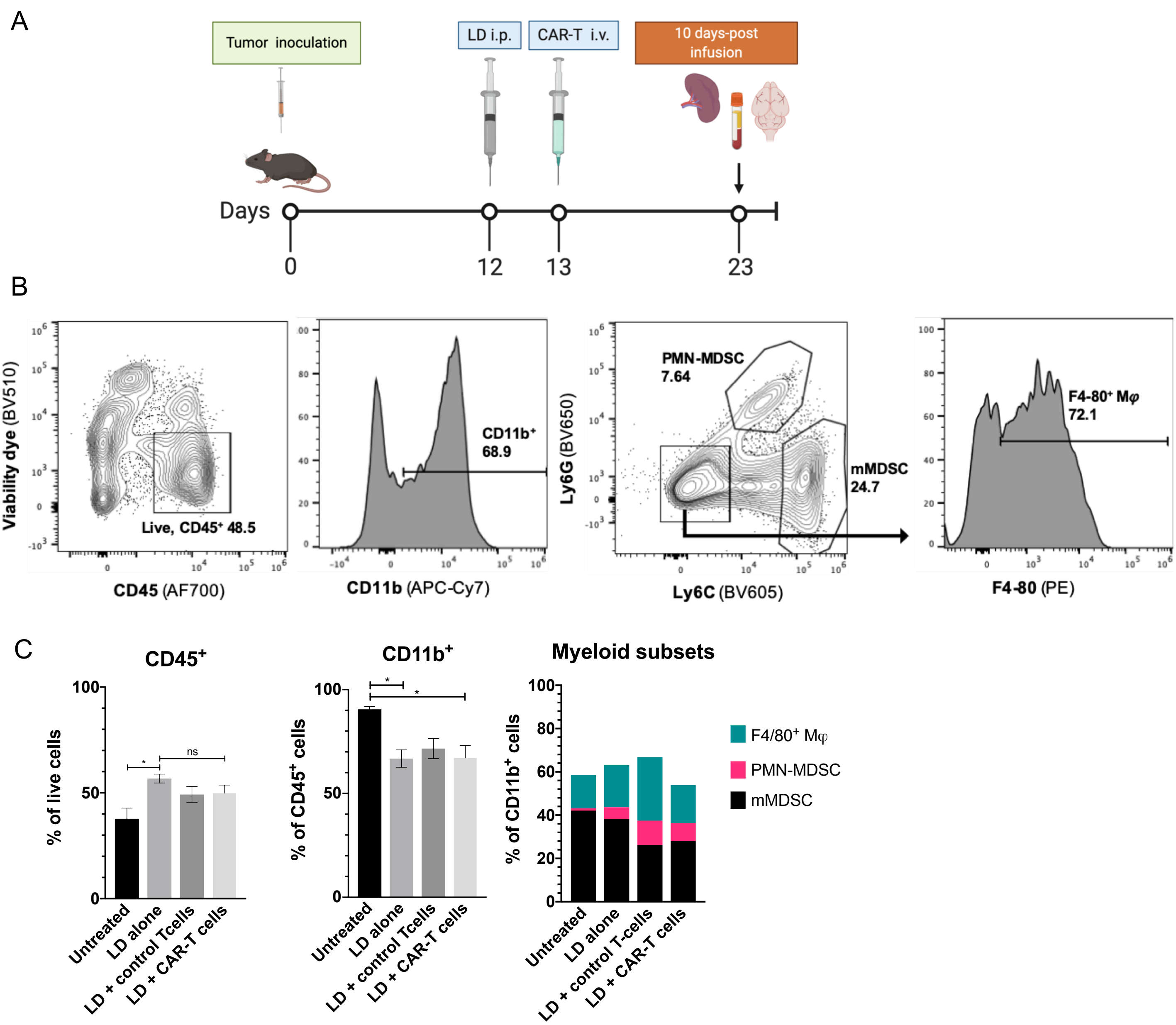

**Suppl. Figure 8. Prospective analysis of BILs 10 days post CART infusion shows no changes in myeloid cell compartment. (A)** Mice (n=12) received intracranial SB28-EGFRvIII tumor cell injections as on day 0. On day 12, mice were randomized to receive no treatment (untreated, n=3) or LD (n=9). On Day 13, LD-treated animals received I.V. infusions of either PBS (LD alone, n=3), $3x10^6$ untransduced T-cells (control T-cells, n=3), or $3x10^6$ CAR-T-cells (LD+CAR-T cells, n=3). Ten days later, animals were sacrificed, and brain-infiltrating leukocytes (BILs) were collected by Percoll gradient centrifugation. BILs were stained and analyzed by flow cytometry for markers characterizing the immune microenvironment. **(B)** Profiling myeloid immune populations in $EGFRvIII^+$ tumor-bearing mice. Equal number of total cells were stained and analyzed. Example flow gating for the different populations shown in **(C).** All samples were gated on single, live events. **(C)** The proportions of different suppressive $CD11b^+$ myeloid cell populations do not change significantly after treatment with anti-EGFRvIII mCAR. Bars represent the mean of 3 biological replicates.

### Supplemental Figure 9

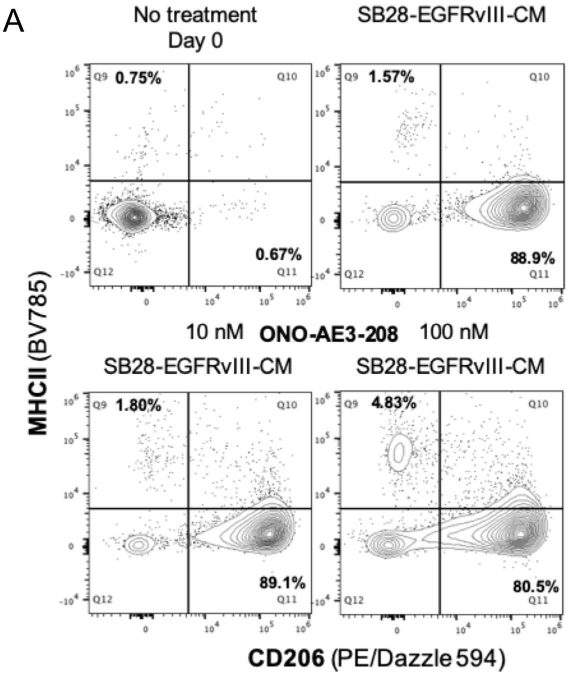


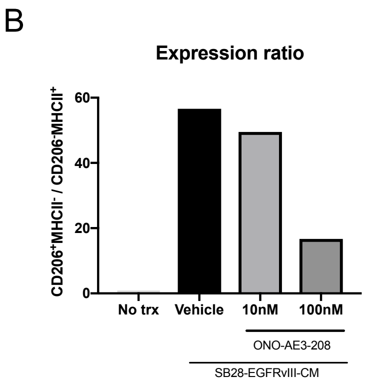

**Suppl. Figure 9. Treatment with ONO-AE3-208 reduces the abundance of pro-tumor markers in CM-treated myeloid cell *in vitro*. (A)** Flow cytometry analysis of CD206 and MHCII expression by SB28-EGFRvIII-CM-treated CD11b$^{+}$ cells. **(B)** Quantification of the ratio of CD206$^{+}$MHCII$^{-}$ (pro-tumor phenotype) to CD206$^{-}$MHCII$^{+}$ (anti-tumor phenotype) cells present in each culture condition shown in **(A)**.

**Supplemental Figure 10**

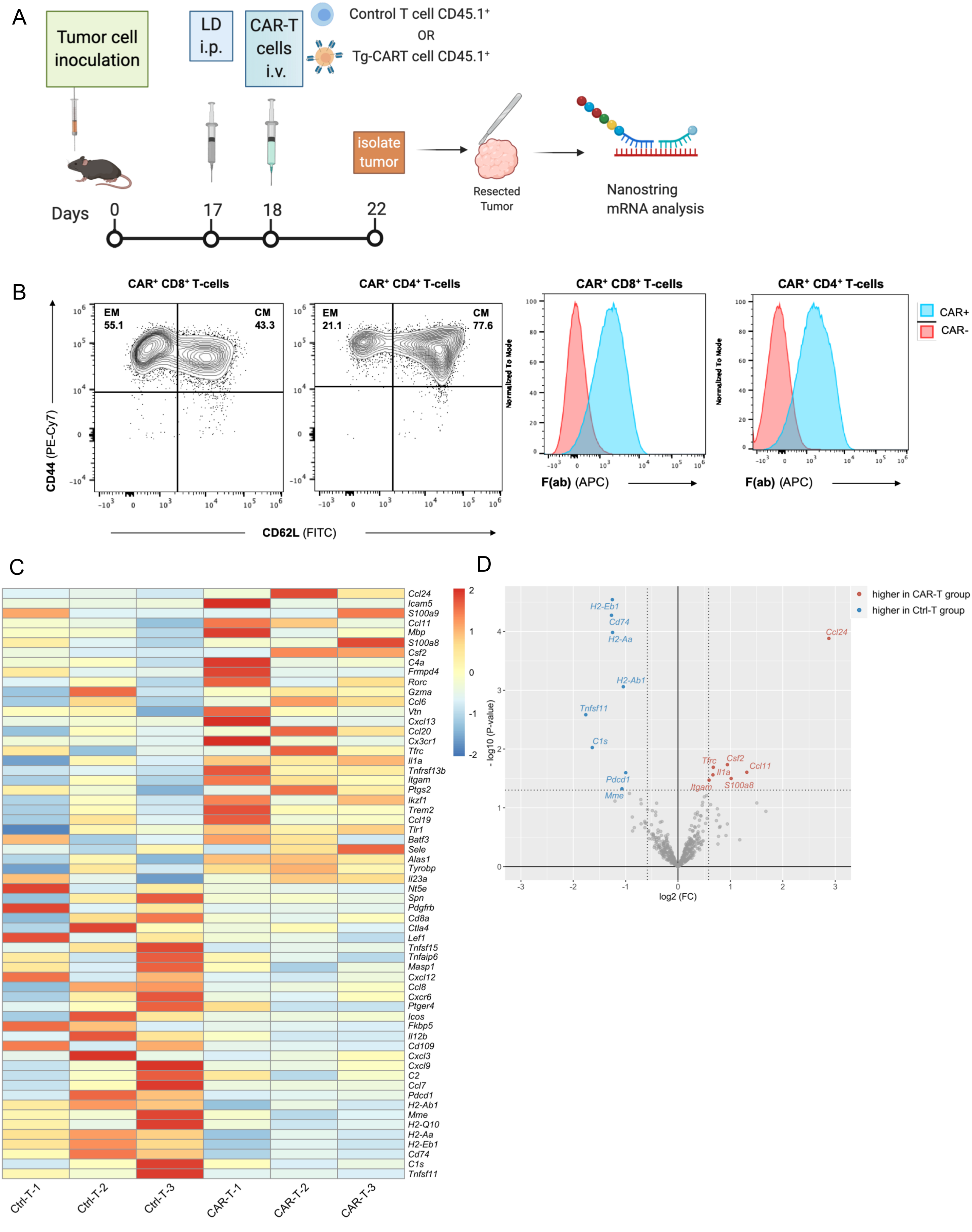

**Suppl. Figure 10. Prospective analysis of tumors 4 days post CART infusion reveals a heterogenous response to therapy. (A)** Mice (n=6) received intracranial SB28-EGFRvIII tumor cell injections as on day 0. On day 17, all mice received a LD regimen treatment. On the following day, mice were randomized and received an i.v. infusion of either $3x10^6$ control T-cells (Ctrl-T, n=3) or $3x10^6$ Tg-CART-cells (CAR-T, n=3). Four days later, animals were sacrificed, and tumor mRNA samples were analyzed using the Mouse Immunology v2 panel by Nanostring. **(B)** Infusion product administered by i.v. ACT in **(A)** was analyzed by FC for surface T-cell memory markers and the expression of the mCAR by F(ab) staining. **(C)** Heatmap showing the expression status of the top and bottom 30 CART-cell regulated genes according to FC values determined by DE-analysis. The colors indicate the row-scaled expression levels. **(D)** Volcano plot summarizing the differential expression (DE) analysis of tumor-bearing mice treated with control-T cells (Cntl-T) or Tg-CART-cells [(CAR-T), n=3 each]. X- and Y- axes indicate log2-scaled fold changes (FC) and minus log10-scaled unadjusted p-values, respectively. The differentially expressed genes were highlighted in color and labeled based on the cut-off values of |FC|>1.5 and p-value<0.05 (dotted line). Detailed gene expression values for highlighted genes are shown in **Supplemental Tables 4-5**.
